# DIO3 coordinates photoreceptor development timing and fate stability in human retinal organoids

**DOI:** 10.1101/2025.03.20.644422

**Authors:** Christina McNerney, Clayton P. Santiago, Kiara C. Eldred, Ian Glass, Tom A. Reh, Arturo Hernandez, Seth Blackshaw, Nathan D. Lord, Robert J. Johnston

## Abstract

The mechanisms governing the generation of neuronal subtypes at distinct times and proportions during human retinal development are poorly understood. While thyroid hormone (TH) signaling specifies cone photoreceptor subtypes, how this regulation changes over time remains unclear. To address this question, we studied the expression and function of type 3 iodothyronine deiodinase (DIO3), an enzyme that degrades TH, in human retinal organoids. We show that DIO3 is a master regulator of human photoreceptor developmental timing and cell fate stability. DIO3 is highly expressed in retinal progenitor cells (RPCs) and decreases as these cells asynchronously differentiate into neurons, progressively reducing TH degradation and increasing TH signaling. *DIO3* mutant organoids display precocious development of S cones, L/M cones, and rods, increased photoreceptor (PR) density, and adoption of L/M cone fate characteristics by S cones and rods. Our multiomics and chimeric organoid experiments show that cell autonomous and non-autonomous mechanisms locally coordinate and maintain DIO3 expression and TH signaling levels among cells. Computational modeling reveals a mechanism that couples TH levels and fate specification, providing robustness to photoreceptor development as compared to a probabilistic, cell-intrinsic mechanism. Based on our findings, we propose an ‘hourglass hypothesis’, in which the proportion of progenitors to neurons decreases over time to relieve TH degradation, which triggers development of PR subtypes at specific times. Our study identifies how local regulation of thyroid hormone signaling influences neural cell fate specification, which may be a consideration for designing regenerative therapies.

## Introduction

The cell types of the developing retina are generated in overlapping temporal windows. Lineage tracing studies suggest that the timing of retinal cell type generation is regulated by probabilistic, cell-intrinsic mechanisms and/or extrinsic signaling (Alexiades and Cepko, 1997; Belliveau and Cepko, 1999; Cepko, 2014; Gomes et al., 2011; He et al., 2012; Sakagami et al., 2009). Whereas cell-intrinsic mechanisms involving transcriptional regulation have been extensively studied, our understanding of how signaling controls retinal cell developmental timing is limited (Cepko, 2014; Viets et al., 2016; Wang et al., 2005; Zhang et al., 2023). Here, we investigated how signaling is regulated to control photoreceptor developmental timing in human retinal organoids.

Photoreceptors (PRs) serve essential functions in vision. Rod PRs express Rhodopsin (Rho) and enable low-light vision. Cone PRs provide high-acuity daytime and trichromatic color vision in humans. The three subtypes of cone PRs are defined by the visual pigment that they express: blue-opsin (short wavelength; S), green-opsin (medium wavelength; M), or red-opsin (long wavelength; L). Cone subtypes are diversified by two fate decisions: (1) S vs L/M fates, and (2) L vs M fates (Nathans et al., 1986). In the developing human retina, S-opsin is expressed before L/M-opsin and Rho (Curcio et al., 1990; Xiao and Hendrickson, 2000). Here, we investigated the developmental timing and fate specification of human S cones, L/M cones, and rods.

Thyroid hormone (TH) signaling regulates cone subtype specification and development across species (Aramaki et al., 2022; Lu et al., 2009; Ma et al., 2014; Mackin et al., 2019; McNerney and Johnston, 2021; Ng et al., 2009; Ng et al., 2001; Ng et al., 2017; Ng et al., 2011; Ng et al., 2010; Roberts et al., 2006). We previously showed that terminal S and L/M cone fates are regulated by TH signaling in human retinal organoids (Eldred et al., 2018). Thyroid hormone receptor B (THRB) is a nuclear hormone receptor that mediates TH signaling. *THRBΔ* mutant organoids display a loss of L/M cones, suggesting that THRB is required for red/green cone fate. TH has two forms: the less active, circulating T4, and the highly active T3. *wildtype* organoids grown in high T3 conditions have a high density of L/M cones and a low density of S cones on day 200. When *wildtype* organoids are exposed to high T3 conditions after the generation of S-opsin-expressing cones, these same cells begin to express L/M-opsin, suggesting a cell fate conversion (Hussey et al., 2024). While TH signaling is critical for fate specification, how the retina regulates TH signaling to control the timing of cone subtype development remains unclear.

TH regulation occurs on both organismal and local scales. At the organismal scale, the hypothalamus-anterior pituitary-thyroid (HPT) axis controls TH levels. The hypothalamus secretes thyrotropin-releasing hormone (TRH), which promotes the anterior pituitary to release thyroid stimulating hormone (TSH). TSH acts on the thyroid gland to promote production and release of TH. Circulating TH then regulates the hypothalamus and anterior pituitary in a negative feedback loop: low TH leads to increase TH production, while high TH suppresses TH production (Alkemade et al., 2005; Dietrich et al., 2012; Fekete and Lechan, 2014; Feldt-Rasmussen et al., 2021; Fliers et al., 2006; Gelfand et al., 1987; Larsen, 1982). On the local scale, additional regulation occurs in the thyroid and target tissues, such as the liver, which are capable of activating circulating T4 into T3 or degrading TH (both T3 and T4) to modulate TH signaling (Bianco et al., 2002; Galton, 2017; Gereben et al., 2008a; Muller et al., 2014). These processes are mediated by deiodinase enzymes, which deiodinate THs via a selenocysteine-containing active site to either activate (DIO1, DIO2) or degrade TH (DIO1, DIO3) (Gereben et al., 2008a; Gereben et al., 2008b; Hernandez et al., 2002).

In mice, *Dio3* is expressed in the early developing retina and decreases over time (Ng et al., 2017; Ng et al., 2010). *Dio3Δ* mutant mice display increased cone death, suggesting that elevated TH levels are detrimental to PR and retinal development in mouse (Lu et al., 2009; Ma et al., 2014; Ng et al., 2009; Ng et al., 2017; Ng et al., 2011; Ng et al., 2010; Roberts et al., 2006). In the developing chick retina, *DIO3* is expressed in two waves before decreasing over time (Trimarchi et al., 2008). Similarly, we previously showed that in human fetal retinas and retinal organoids, *DIO3* mRNA is highly expressed early, then gradually decreases as development progresses (Eldred et al., 2018).

The dynamic expression of DIO3, its role in TH degradation, and the influence of TH signaling on cone subtype specification suggest that DIO3 regulates the timing of human photoreceptor development. However, this function has not been explored, partly due to the challenges of obtaining fetal tissue. Stem cell-derived human retinal organoids provide a genetically and pharmacologically tractable model to interrogate mechanisms of human development. Previous studies characterized extensive similarities between fetal retina and organoid development, establishing the utility of organoids for investigating cell fate specification and developmental timing (Eiraku et al., 2011; Eldred et al., 2018; Hadyniak et al., 2024; Hussey et al., 2024; Kaewkhaw et al., 2015; Nakano et al., 2012; Sridhar et al., 2020; Wahlin et al., 2017). In this study, we sought to determine how regulation of TH signaling and timing of PR development are linked during human retinal organoid development. As *DIO3* expression decreases during human retinal and organoid development (Eldred et al., 2018), we hypothesized that loss of TH degradation increases TH signaling, driving PR development over time.

To test this hypothesis, we examined how DIO3 controls the temporality of PR development. DIO3 is expressed in retinal progenitor cells (RPCs) and its expression and activity decrease as RPCs differentiate. *DIO3* mutant human organoids display precocious cone and rod development, increased PR density, and non-exclusive PR fates. Importantly, both cell autonomous and non-autonomous mechanisms regulate DIO3 levels to coordinate TH signaling and PR development locally in the retina. Thus, as the RPC:neuron proportion decreases over time, TH degradation is relieved, progressively increasing TH signaling to trigger PR subtype development.

## Results

### DIO3 is expressed in RPCs and decreases in neurons in fetal retinas and organoids

To determine how *DIO3* expression changes during human retinal development, we examined a published single nucleus RNA-seq (snRNA-seq) dataset of the developing human fetal retina (Zuo et al., 2024) and found that *DIO3* mRNA is highly expressed in the RPC cluster (**Fig. 1A-B**), identified by co-expression of *VSX2* and *PAX6* (**Fig. S1A-C**). We also observed DIO3 protein expression in VSX2*+* RPCs in a 14-week old fetal retina (**Fig. 1C**). In the snRNA-seq dataset, minimal *DIO3* mRNA expression was observed in mature neurons (**Fig. 1A-B**). These data suggest that DIO3 is expressed in RPCs and decreases as RPCs differentiate into terminal retinal cells.

**Fig. 1.**
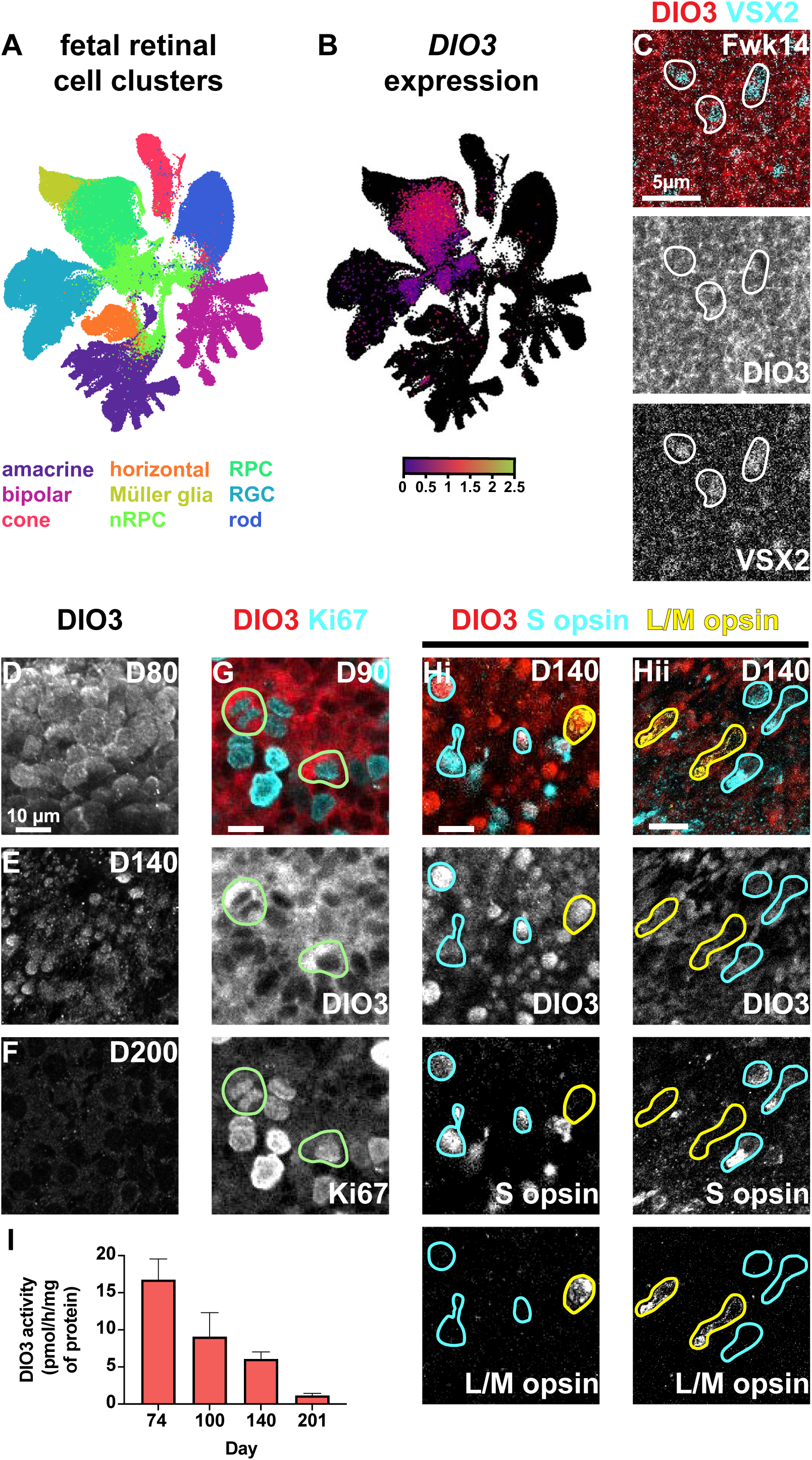
DIO3 is expressed in RPCs in the developing human retina and retinal organoids. (A) UMAP from CZ CELLxGENE Discover (Zuo et al., 2024) of human fetal retina scRNA-seq with major cell types labeled. (B) *DIO3* mRNA expression in fetal retina limited to DIO3+ cells, overlaid on the unannotated UMAP. (C) DIO3 is expressed in VSX2+ cells in a 14 week old fetal retina. (D-F) DIO3 expression in human retinal organoids at (D) day 80, (E) 140, and (F) 200. (G) DIO3 and Ki67 expression in human retinal organoids at day 90. Green circles indicate Ki67+/DIO3+ cells. (H) DIO3 and cone opsin expression in human retinal organoids at day 140. Yellow outlines indicate L/M-opsin+ cells. Blue outlines indicate S-opsin+ cells. (Hi) Region of morphologically naïve photoreceptors within (H) organoid. (Hii) Region of morphologically more mature photoreceptors within (H) organoid. (I) DIO3 activity as measured by deiodination activity of *wildtype* organoids over time. Error bars represent SEM.

As accessibility and experimental tractability of human fetal retinal tissue are limited, we studied human retinal organoids to evaluate the dynamics of DIO3 expression. We visualized DIO3 protein on days 80, 90, 140, and 200 of organoid development (**Fig. 1D-H**). On days 80 and 90, DIO3 was highly expressed in RPCs (**Fig. 1D, G**). On day 140, DIO3 was highly expressed in a subset of cells but was low in others (**Fig. 1E**). On day 200, DIO3 was minimal across all cells (**Fig. 1F**). These data suggest that the proportion of retinal cells expressing DIO3 decreases during development.

We next related cell-type-specificity to temporal dynamics of DIO3 expression. In organoids on day 90, DIO3 was observed in cells that express Ki67 (**Fig. 1G**), which marks mitotic cells. This observation suggests that DIO3 is expressed in mitotic RPCs in organoids, similar to *DIO3* mRNA in fetal retinas (**Fig. 1A-B**). To determine the relationship between DIO3 and cone differentiation, we examined expression of DIO3 in cells that express S-opsin or L/M-opsin. We determined maturity of these cells by comparing morphological characteristics of each cell within the same organoid. Whereas immature cones had round cell bodies with an even distribution of opsin throughout the cell, mature cones had elongated processes and opsin localized to putative inner segments. In organoids on day 140, DIO3 was expressed at higher levels in immature S-opsin expressing cones and L/M-opsin expressing cones (**Fig. 1Hi**) and lower levels in mature S-opsin expressing cones and L/M-opsin expressing cones (**Fig. 1Hii**). Finally, to relate DIO3 expression and function, we measured DIO3 deiodination activity in retinal organoids and found that it decreases during development (**Fig. 1I**), consistent with the progressive reduction in DIO3 protein expression.

These results suggest that DIO3 is expressed in RPCs and decreases as RPCs differentiate into neurons. As the proportion of DIO3-expressing cells decreases, DIO3 activity decreases.

### Generation of DIO3 mutant organoids

To investigate the functional role of DIO3 in human retinal organoids, we used CRISPR/Cas9 to generate a stem cell line with a homozygous 4 base pair deletion that introduced an early stop, yielding a presumptive null mutation (“*DIO3^0^”,* **Fig. S1D-E**). We differentiated *DIO3^0^* null mutant organoids, which did not generate retinal tissue as indicated by the absence of optic cup formation and lamination (0 of 384 aggregates developed retinal tissue across 2 independent differentiations), while *wildtype* control organoids successfully differentiated in parallel (**Fig. S1F-H**). This suggests that *DIO3* is required for retinal tissue development in organoids.

To circumvent this requirement, we generated the *DIO3^Δ33^/+* stem cell line, which is heterozygous for a wildtype allele and a 33 base pair, in-frame deletion (“*DIO3^Δ33^*”, **Fig. S1I**), causing a loss of 11 amino acids adjacent to, but not including the active site (**Fig. S1J**). We differentiated *DIO3^Δ33^/+* organoids, which developed normal optic cup-like morphologies and lamination (**Fig. S1K**). In *DIO3^Δ33^/+* organoids, DIO3 protein remained detectable by IHC on day 100 (**Fig. 2A**), consistent with the generation of DIO3 protein that retained the antigenic epitope (epitope positions 250-300, **Fig. S1J**).

**Fig. 2.**
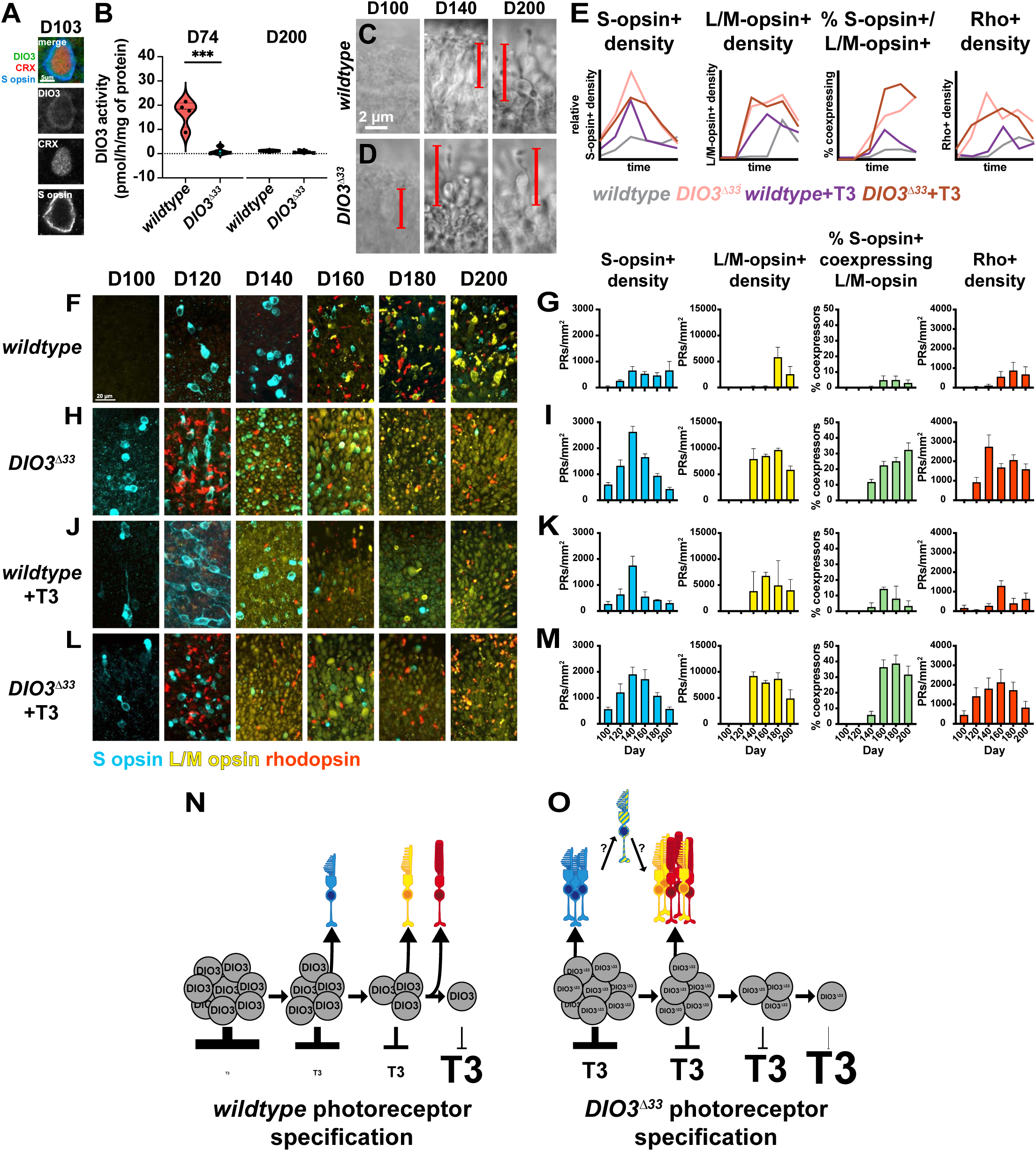
*DIO3^Δ33^* mutant and +T3 conditions accelerate PR development, increase PR quantities, and yield mixed PR fates. (A) DIO3 expression in an S-opsin+, CRX+ cell day 103 *DIO3^Δ33^* organoid cell. (B) DIO3 activity of *wildtype* and *DIO3^Δ33^* at day 74 and 201 of differentiation. At day 74, p=0.0004; at day 201, p=0.2225 (ns) by unpaired t test. (C-D) Brightfield imaging of *wildtype* (C) and *DIO3^Δ33^* (D) organoids at day 100 (left), 140 (center), and 200 (right). Bracketed lines highlight individual PR precursors or PRs. (E) Graphical representation of (G, I, K, M) quantifications. Y axes are scaled relative to each opsin. (F, H, J, L) Timeline of opsin expression in human retinal organoids at day 100, 120, 140, 160, 180, and 200 in (F) *wildtype* organoids, (H) *DIO3^Δ33^*mutant retinal organoids, (J) *wildtype* +T3 organoids, and (L) *DIO3^Δ33^*+T3 organoids. (G, I, K, M) Quantification of S-opsin+ cell density (blue), L/M-opsin+ cell density (yellow), %L/M-opsin+ cells expressing S-opsin (green), Rho+ cell density (red) for (G) *wildtype* organoids, (I) *DIO3^Δ33^* mutant retinal organoids, (K) +T3 treated *wildtype* organoids, and (M) *DIO3^Δ33^*+T3 organoids. Error bars represent SEM. *Wildtype*: D100 N=5, D120 N=8, D140 N=4, D160 N=4, D180 N=6, D200 N=4. *DIO3^Δ33^*: D100 N=5, D120 N=9, D140 N=9, D160 N=10, D180 N=17, D200 N=11. *wildtype* +T3: D100 N=5, D120 N=3, D140 N=3, D160 N=3, D180 N=2, D200 N=4. *DIO3^Δ33^* +T3: D100 N=4, D120 N=3, D140 N=4, D160 N=4, D180 N=3, D200 N=4. Statistics reported in Fig. S2. (N) Model of *wildtype* photoreceptor specification, in which DIO3-expressing RPC populations decrease and T3 levels increase during development. (O) Model of *DIO3^Δ33^*photoreceptor specification, in which the *DIO3^Δ33^*-expressing RPC population has impaired T3 degradation, leading to high T3 levels. S-opsin+ cells are generated early and likely convert to L/M cones. L/M cones and rods are generated early and at higher densities than in *wildtype*, with some rods co-expressing L/M opsin.

To assess changes in DIO3 enzymatic activity in *DIO3^Δ33^/+* organoids, we evaluated deiodination activity in *wildtype* and *DIO3^Δ33^/+* organoids on day 74, when *DIO3* mRNA and protein are highly expressed (**Fig. 1D, G**) (Eldred et al., 2018). Deiodination activity in *DIO3^Δ33^/+* was reduced by ∼94% compared to *wildtype* organoids (*wildtype* = 16.8 pmol/h/mg of protein, *DIO3^Δ33^/+* = 1.0 pmol/h/mg of protein, P<0.001) (**Fig. 2B**). Since the *DIO3^Δ33^/+* organoids are heterozygous, this >50% reduction in deiodination activity suggests that the *DIO3^Δ33^/+* mutation has a dominant negative phenotype. We observed a similar decrease in DIO3 function in 76-day-old *wildtype* and *DIO3^Δ33^/+* organoids derived from an independent differentiation (*wildtype* = 17.4 pmol/h/mg of protein, *DIO3^Δ33^/+* = 1.0 pmol/h/mg of protein, P<0.01) (**Fig. S1L**). We also measured deiodination activity in *wildtype* and *DIO3^Δ33^/+* organoids on day 200 and observed low activity in both (*wildtype* = 1.2 pmol/h/mg of protein, *DIO3^Δ33^/+* = 0.7 pmol/h/mg of protein, ns) (**Fig. 2B**), consistent with the reduction in DIO3 activity and expression in *wildtype* organoids at this late timepoint (**Fig. 1F, I**) and the decreased deiodination function in *DIO3^Δ33^/+.* Notably, the remaining, minimal DIO3 activity permits retinal tissue generation, allowing for characterization of mutant phenotypes. We refer to this stem cell line as “*DIO3^Δ33^*mutant” for simplicity.

### DIO3^Δ33^ mutants display changes in PR developmental timing, quantity, and stability

To determine the roles of DIO3 and T3 in PR development, we differentiated organoids in parallel in four conditions: *wildtype, DIO3^Δ33^*, *wildtype +*T3, and *DIO3^Δ33^ +*T3. We evaluated PR development based on the densities of S-opsin-expressing cells (S-opsin+ cells), L/M-opsin-expressing cells (L/M-opsin+ cells), and Rho-expressing cells (Rho+ cells) on days 100, 120, 140, 160, 180, and 200. We also observed and quantified cells that co-expressed S-opsin and L/M-opsin (% S-opsin+ co-expressing L/M-opsin) and Rho and L/M-opsin (% Rho+ co-expressing L/M-opsin).

To assess how DIO3 regulates PR developmental timing, we imaged *wildtype* and *DIO3^Δ33^*mutant organoids with brightfield microscopy (**Fig. 2C, D**). At day 100, *wildtype* organoids had no visible PRs (**D100, Fig. 2C**), whereas *DIO3^Δ33^* organoids had immature PRs with elongated, goblet-shaped cell bodies (**D100, Fig. 2D**). At day 140, *wildtype* organoids had a low density of immature PRs (**D140, Fig. 2C**), while *DIO3^Δ33^* organoids had mature PRs with outer segments (**D140, Fig. 2D**). At day 200, both *wildtype* (**D200, Fig. 2C**) and *DIO3^Δ33^* organoids (**D200, Fig. 2D**) had dense PRs with mature morphologies. These observations suggest that PR development is accelerated in *DIO3^Δ33^*mutant organoids.

To investigate PR subtype development in *DIO3^Δ33^* mutants, we next compared developmental timing of S-opsin+, L/M-opsin+, and Rho+ cells in *wildtype* and *DIO3^Δ33^* mutant organoids with IHC of opsins (**Fig. 2E-I**, statistics reported in **Fig. S2**). At day 100, *wildtype* organoids contained very few S-opsin+ cells (**D100, Fig. 2E-G, Fig. S2A**). In contrast, *DIO3^Δ33^* organoids displayed significantly more S-opsin+ cells (S-opsin+, *wildtype* = 39 PRs/mm^2^, *DIO3^Δ33^* = 602 PRs/mm^2^, P<0.001) (**D100, Fig. 2E, 2H-I, Fig. S2A**), suggesting that development of S-opsin+ cells is accelerated in *DIO3^Δ33^*mutants.

At day 120, *wildtype* organoids contained morphologically immature S-opsin+ cells, (**D120, Fig. 2E-G**), whereas *DIO3^Δ33^*organoids displayed an increase in S-opsin+ cells, which localized to the outer nuclear layer (ONL) and had inner segments, indicating maturity (S-opsin+, *wildtype* = 259 PRs/mm^2^, *DIO3^Δ33^* = 1325 PRs/mm^2^, P<0.01) (**D120, Fig. 2E, 2H-I, Fig. S2A**). *DIO3^Δ33^* organoids also contained a small population of L/M-opsin+ cells (L/M-opsin+, *wildtype* = 0 PRs/mm^2^, *DIO3^Δ33^* = 12 PRs/mm^2^, ns) and a larger population of Rho+ cells (Rho+, *wildtype* = 42 PRs/mm^2^, *DIO3^Δ33^* = 925 PRs/mm^2^, P<0.05) (**D120, Fig. 2E, 2H-I, Fig. S2B, D),** suggesting that development of L/M-opsin+ and Rho+ cells is also accelerated in *DIO3^Δ33^* mutants.

At day 140, *wildtype* organoids exhibited increased densities of S-opsin+ cells and few L/M opsin+ and Rho+ cells (**D140, Fig. 2E-G, Fig. S2A, B, D**). In contrast, *DIO3^Δ33^* organoids displayed dramatic increases in S-opsin+, L/M-opsin+, and Rho+ cells (S-opsin+, *wildtype* = 653 PRs/mm^2^, *DIO3^Δ33^*= 2632 PRs/mm^2^, P<0.001; L/M-opsin+, *wildtype* = 190 PRs/mm^2^, *DIO3^Δ33^* = 7936 PRs/mm^2^, ns; Rho+, *wildtype* = 99 PRs/mm^2^, *DIO3^Δ33^* = 2746 PRs/mm^2^, P<0.05) (**D140, Fig. 2E, 2H-I, Fig. S2A, B, D**). A subset of L/M-opsin+ cells had simpler morphologies and lower L/M-opsin expression (**D140, Fig. 2H**). These L/M-opsin+ cells expressed ARR3 (cone gene, **Fig. S3A-D**), CRX (PR precursor and PR gene, **Fig. S3A-D**) and RCVRN (PR gene, **Fig. S3E-H**), but not NRL (rod gene, **Fig. S3E-H**), confirming their identity as L/M cones. These cells developed outer segments over time (**Fig. SJ**). The length of the outer segments (**Fig. S3K**) and the proportion of lowly expressing L/M-opsin+ cells with outer segments (**Fig. S3L**) increased more rapidly in *DIO3^Δ33^* mutants compared to *wildtype* organoids.

The densities of all PR types were greater in *DIO3^Δ33^*mutant organoids at D140 than at any timepoint in *wildtype* organoid development, suggesting that PR generation is increased in *DIO3^Δ33^*mutants (**Fig. S2A, B, D)**.

At day 140, *DIO3^Δ33^* organoids developed a population of S-opsin+ cells that co-expressed L/M-opsin (S&L/M-opsin+, *wildtype* = 1%, *DIO3^Δ33^* = 12%, P<0.01) (**D140, Fig. 2E, 2H-I, S3L, Fig. S3I**). We previously showed that co-expression is indicative of conversion of S-opsin+ cells to L/M fate (Hussey et al., 2024). We also observed a population of Rho+ cells that co-expressed L/M-opsin (L/M-opsin&Rho+, *wildtype* = 0%, *DIO3^Δ33^* = 43%, P<0.0001;) (**Fig. S2E, S3M-T**). These results suggest that photoreceptor fate stability is disrupted in *DIO3^Δ33^* organoids.

Across days 160, 180, and 200, *wildtype* organoids maintained S-opsin+ cell density and increased L/M-opsin+ and Rho+ cell densities (**D160, D180, D200 Fig. 2E-G, Fig. S2A, B, D**). In contrast, *DIO3^Δ33^* organoids displayed a dramatic decrease in the density of S-opsin+ cells, an increase in the populations of S-opsin+ or Rho+ cells that co-expressed L/M-opsin, and similar densities of L/M-opsin+ and Rho+ cells (**D160, D180, D200, Fig 2E, 2H-I, Fig. S2**). The decrease in S-opsin+ and Rho+ cells from their peaks on day 140 together with the increase in the percentage of co-expressing cells suggest that subsets of S-opsin+ cells and Rho+ cells are converted to L/M cone fate.

Together, *DIO3^Δ33^* organoids displayed accelerated and increased generation of S-opsin+, L/M-opsin+, and Rho+ cells and non-exclusive opsin expression (**Fig 2E, 2N-O, Fig. S2, Fig. S3M-T**), suggesting that DIO3 functions to control the timing, quantity, and stability of PR development.

### Organoids grown in high T3 conditions display similar developmental changes as DIO3^Δ33^ mutant organoids

Since DIO3 degrades T3, *DIO3^Δ33^* mutant organoids likely have elevated TH signaling. Therefore, *wildtype* organoids grown in high exogenous T3 conditions (*wildtype* +T3) should show similar changes in PR timing, quantities, and fate stability as *DIO3^Δ33^* mutant organoids. In *wildtype* organoids grown in high T3 conditions S-opsin+, L/M-opsin+, and Rho+ cells were generated earlier and in higher quantities (**Fig. 2E, 2J-K, Fig. S2A, B, D**). Over time, S-opsin+ cells decreased, while L/M-opsin+ cells and the proportion of S-opsin+ cells co-expressing L/M-opsin increased (**Fig. 2E, 2J-K, Fig. S2A, B, C**). The proportion of Rho+ cells co-expressing L/M-opsin increased at D160 but decreased over the rest of development (**Fig. S2E, Fig. S3P, Q, S**). *wildtype* +T3 organoids displayed less severe changes compared to *DIO3^Δ33^* organoids, likely due to buffering by endogenous, functional DIO3. These observations confirm that the developmental changes observed in *DIO3^Δ33^* mutant organoids are driven by increased TH signaling.

We next tested how addition of T3 affected *DIO3^Δ33^* mutant organoids. *DIO3^Δ33^* mutant organoids grown in high T3 conditions from day 42 to day 200 (*DIO3^Δ33^*+T3) displayed similar phenotypes as *DIO3^Δ33^* mutant organoids alone (**Fig. 2E, 2L-M, Fig. S2, Fig. S3P, Q, T**), consistent with both perturbations affecting the same pathway.

Together, these data suggest that the phenotypes observed in *DIO3^Δ33^* mutant organoids are due to elevated T3 signaling.

### DIO3^Δ33^ mutant PRs are more mature than wildtype PRs on day 200

Our analysis of *DIO3^Δ33^* mutant organoids suggested that PR development is accelerated. To determine if *DIO3^Δ33^* mutant PRs are more mature than *wildtype* PRs, we conducted snRNA-seq and snATAC-seq multi-omics of *wildtype* and *DIO3^Δ33^* organoids on day 200. Clustering based on snRNA-seq alone, snATAC-seq alone, or both identified neural retinal cells including RPCs and all major classes of retinal cells (i.e. cones, rods, amacrine cells, horizontal cells, bipolar cells, Müller glia), except for RGCs, which are lost in organoids by day 200 (**Fig. 3A, S4A-E)** (Cowan et al., 2020; Eldred and Reh, 2021; Sridhar et al., 2020). We also identified non-neural retinal cells including RPE cells, glia-like cells, astrocytes, choroid plexus cells, as well as non-retinal brain and spinal cord-like (BSL) cells, which we previously found in transplanted organoids using scRNA-seq analysis (**Fig. 3A, S4A-D**) (Liu et al., 2023).

**Fig. 3.**
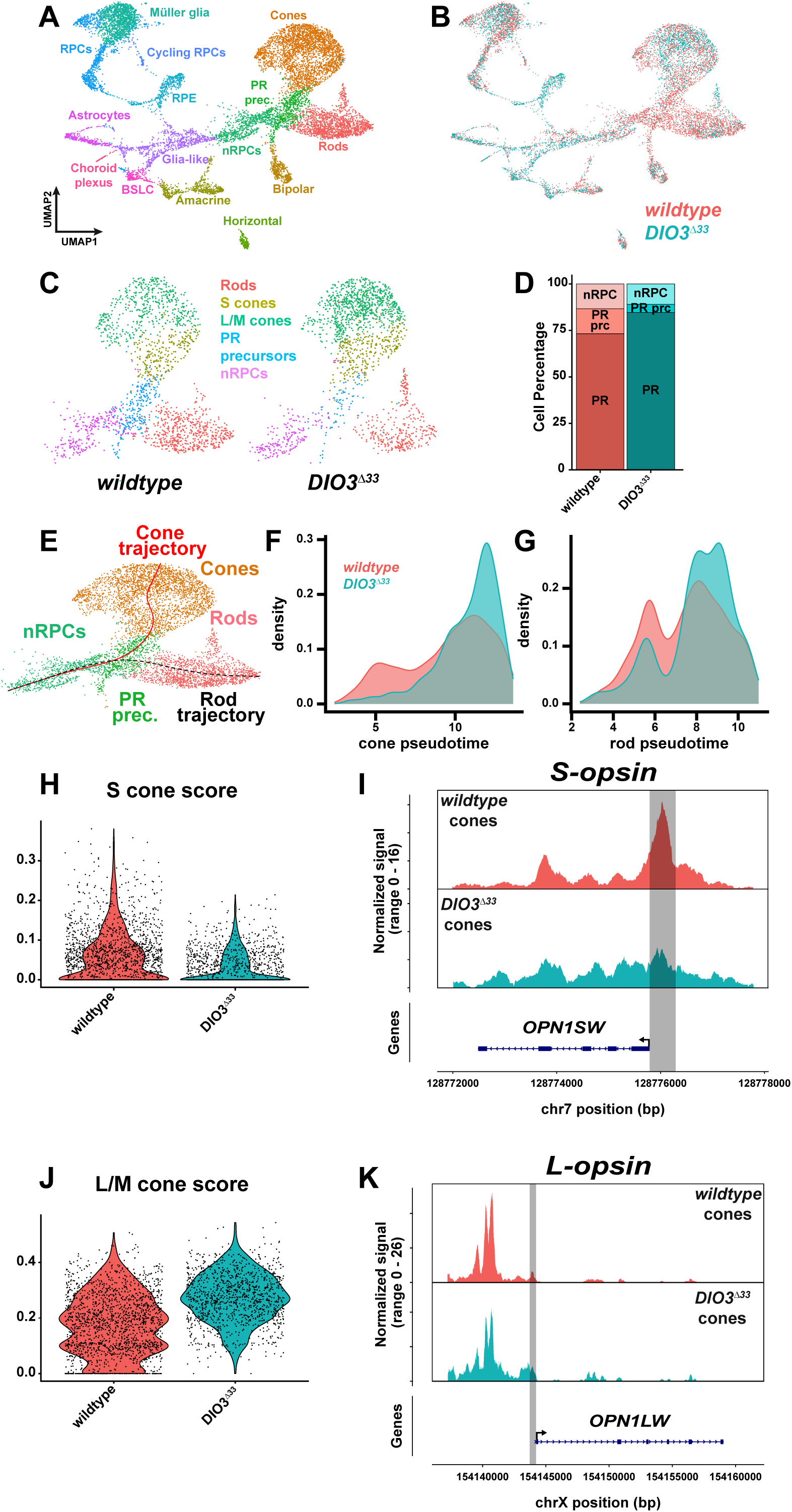
Single nucleus multiomics analysis suggests increased PR maturity and enrichment of L/M cone fate in *DIO3^Δ33^* organoids. (A) Combined UMAP of D200 *DIO3^Δ33^* and *wildtype* organoids. Clusters annotated and color-matched with cell types. (B) Cells colored by tissue origin. Coral = *wildtype* controls and teal = *DIO3^Δ33^*. (C) Zoomed in view of photoreceptor lineages from (A, B) and split by genotype. Neurogenic retinal progenitor cells (nRPCs) in purple, photoreceptor (PR) precursors in blue, rods in red, L/M cones in green, S cones in gold. Cell populations downsampled to visually compare proportions, all other comparisons were performed on complete, non-downsampled data. (D) Proportions of cell states in photoreceptor lineages. Neurogenic RPCs = nRPCs, PR precursors = PR prc, and terminal PRs = PR. (E) Predicted developmental trajectories of photoreceptors. (F-G) Pseudotime density plots of cones (F) and rods (G). X-axis represents distance along pseudotime trajectory as identified in (E), excluding nRPCs. (H, J) S cones (H) or L/M cone (J) scores applied to cones in *wildtype* (coral) and *DIO3^Δ33^* (teal). (I, K) Chromatin accessibilities of *S-opsin* (I) and *L-opsin* (K) genes. Top, tracks separated by genotype. Below, gene annotation. Grey regions outline promoters as defined by 500 bp upstream of the start site.

Since clustering based on snRNA-seq, snATAC-seq, or both resulted in similar outcomes (**Fig. S4A-C**), we initially focused on snRNA-seq-based clustering (**Fig. 3A-B**) and on retinal cells (**Fig. S4E**). We identified populations of neurogenic RPCs (nRPCs), postmitotic PR precursors, and terminally differentiated PRs in *wildtype* and *DIO3^Δ33^* organoids (**Fig. 3C, Fig. S4F**). While *wildtype* and *DIO3^Δ33^* organoids had similar proportions of nRPCs (**Fig. 3C-D**), *DIO3^Δ33^* organoids had mature PRs at the expense of PR precursors (**Fig. 3C-D**), consistent with accelerated PR development in *DIO3^Δ33^* mutants.

Given the acceleration of PR development based on opsin expression in *DIO3^Δ33^* mutants, we analyzed the developmental trajectories of PRs using the snRNA-seq data. We mapped developmental trajectories of PRs, starting from nRPCs, progressing to PR precursors, and then splitting into either rod or cone fates (**Fig. 3E**), consistent with previous studies (Brzezinski and Reh, 2015; Lu et al., 2020; Lyu et al., 2021). To assess developmental maturity, we generated pseudotime density plots for both the cone and rod trajectories. In both cases, a larger proportion of *DIO3^Δ33^* cells clustered at more advanced pseudotime states compared to *wildtype* cells (**Fig. 3F-G**), suggesting that cones and rods are more mature in *DIO3^Δ33^* mutants.

### DIO3^Δ33^ mutant PRs are more L/M cone-like

Our analysis of *DIO3^Δ33^* mutant organoids suggested that a subset of S cones take on L/M cone fate character. In scRNA-seq analysis of human retinal organoids, S and L/M cone subtype clusters are not directly identifiable (Hussey et al., 2024), To assess the fates of cones, we analyzed S- and L/M-cone fate characteristics across all annotated cones. We generated S and L/M cone scores by analyzing the differential expression of genes associated with these cone fates in human (Lu et al., 2020), non-human primate (Peng et al., 2019), and cow (Zhang et al., 2019) and applied these scores to each genotype. When visualized on UMAP plots, the S and L/M cone scores (**Fig. S4G, I**), as well as a rod score, (**Fig. S4L**) correlated with subtype-specific gene expression (**Fig. S4H, J, K, M**), validating our annotation strategy. *DIO3^Δ33^* mutant cones had a lower S cone score and a greater L/M cone score compared to *wildtype* cones (**Fig. 3H, J**), suggesting that the *DIO3^Δ33^* mutant cones are more L/M cone-like. In *DIO3^Δ33^*mutant cones, the *S-opsin* locus is less accessible and the *L-opsin* promoter is more accessible compared to *wildtype* cones (**Fig. 3I, K)**, in line with the loss of S-opsin+ cells and generation of L/M+ cells in *DIO3^Δ33^* mutants on day 200 (**Fig. 2E, 2H-I**). These data are consistent with the co-expression of L/M opsin in S-opsin+ cells in *DIO3^Δ33^* mutants.

### DIO3 expression is regulated by feedback

We next sought to determine how DIO3 and TH signaling are regulated within and between retinal cells. On the organismal scale, TH levels are regulated by a negative feedback loop in which excess circulating T4 inhibits generation and release of TH from the thyroid gland (Gelfand et al., 1987; McNerney and Johnston, 2021). Considering the coordination of development across retinal tissue, we hypothesized that TH regulation might involve additional cell-intrinsic or extrinsic feedback mechanisms.

To assess feedback regulation, we analyzed our snRNA-seq data from organoids on day 200. In *wildtype* organoids, *DIO3* mRNA was expressed in RPCs and sparsely expressed in neurons (**Fig. 4A**), consistent with snRNA-seq analysis of human fetal retinal expression (**Fig. 1A-B**) and our assessment of *DIO3* expression in organoids (**Fig. 1D-H**). In *DIO3^Δ33^* organoids, *DIO3* mRNA was upregulated and the *DIO3* locus was more accessible across all cells (**Fig. 4A-D**) and in cones (**Fig. S5A-B)** and rods (**S5C-D**) specifically. Similarly, DIO3 protein was dramatically upregulated in *DIO3^Δ33^* mutant organoids (**Fig. 4E-F**). These data suggest that *DIO3* gene expression is upregulated when DIO3 function is impaired.

**Fig. 4.**
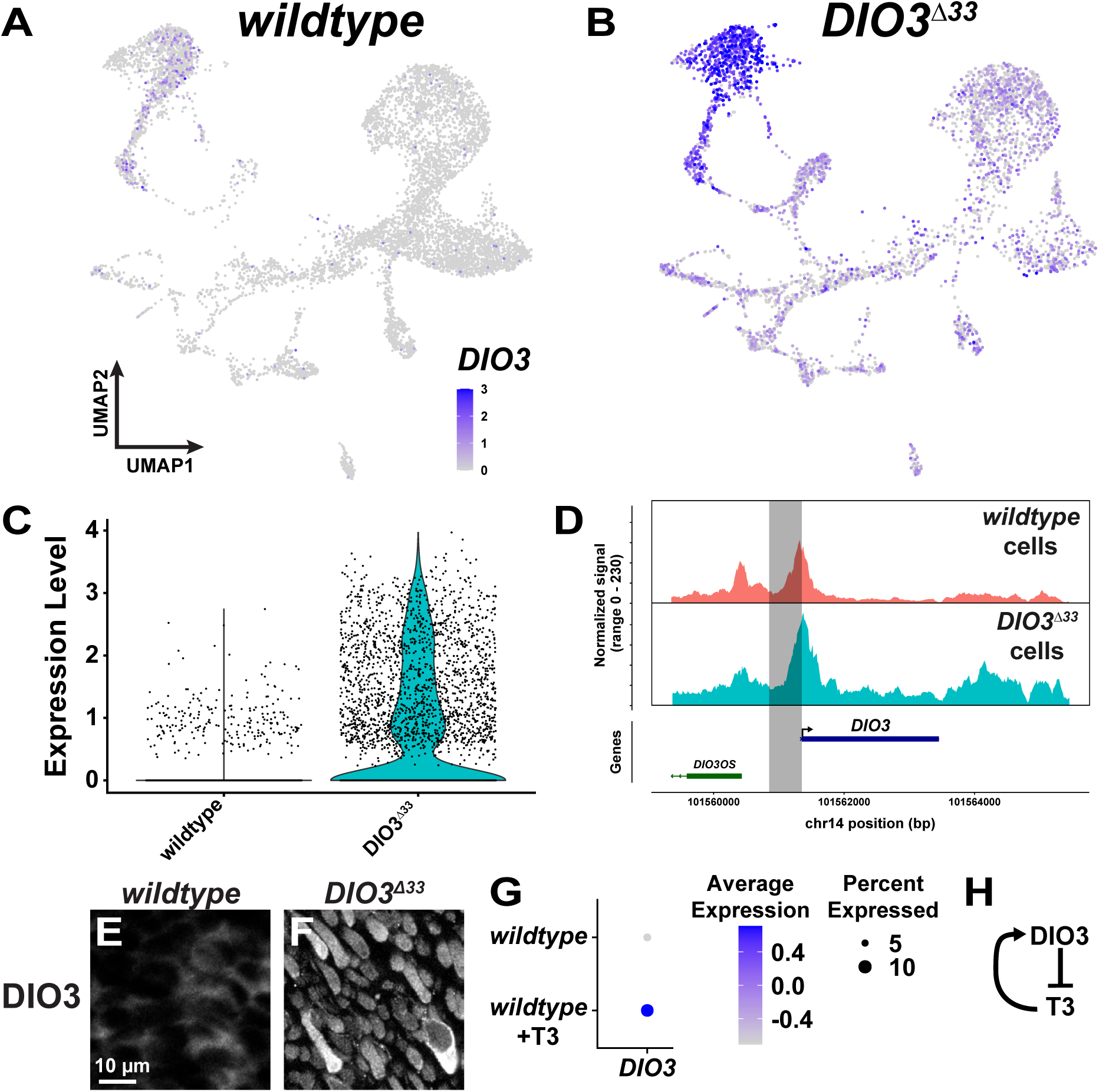
*DIO3* is controlled by feedback. (A, B) Expression of *DIO3* mRNA in *wildtype* (A) and *DIO3^Δ33^* (B) organoids. (C) *DIO3* mRNA expression in retinal cells in *wildtype* and *DIO3^Δ33^* organoids. (D) Top: Accessibility of *DIO3* locus in all retinal cells split by genotype. Bottom: Annotated gene locus (E-F) DIO3 protein expression in *wildtype* (E) or *DIO3^Δ33^* (F) organoids at day 180. The antigenic epitope is intact in *DIO3^Δ33^* mutants and the protein is detectable by IHC (Fig. 2A, Fig. S1B, E) (G) *DIO3* mRNA expression in *wildtype* and *wildtype* +T3 conditions (Hussey et al., 2024). (H) Model of feedback network.

Since DIO3 degrades TH and *DIO3* expression increases in *DIO3^Δ33^*mutant organoids, we predicted that high T3 conditions would also lead to an increase in *DIO3* expression. To test this hypothesis, we analyzed our previously published scRNA-seq data generated from day 200 *wildtype* organoids and *wildtype+T3* organoids (Hussey et al., 2024). *DIO3* expression increased in high T3 conditions (**Fig. 4G**), suggesting that TH signaling activates *DIO3* expression.

Together, these results suggest that DIO3 and TH interact in a feedback loop: DIO3 degrades TH, which in turn activates *DIO3* expression (**Fig. 4H**). This negative feedback mechanism is consistent with local homeostatic regulation of *DIO3* expression and TH signaling.

### DIO3 is regulated by cell non-autonomous feedback

Since retinal development is coordinated and DIO3 regulates TH levels to control timing of PR development, we hypothesized that DIO3 and TH signaling are regulated by intercellular feedback. To test this, we generated two sets of chimeric organoids. We used three types of stem cells: *wildtype* H7 stem cells, *DIO3^Δ33^* mutant H7 stem cells, and *CRX:tdTomato wildtype* H9 stem cells. In the control, we mixed unmarked *wildtype* H7 cells with *CRX:tdTomato wildtype* H9 cells to account for potential differences in hESC line differentiation dynamics. In the experimental set, we mixed *DIO3^Δ33^* H7 mutant cells with *CRX:tdTomato wildtype* H9 cells. From these mixes of stem cells, we differentiated the heterogeneous aggregates into chimeric retinal organoids (**Fig. 5A-B**).

**Fig. 5.**
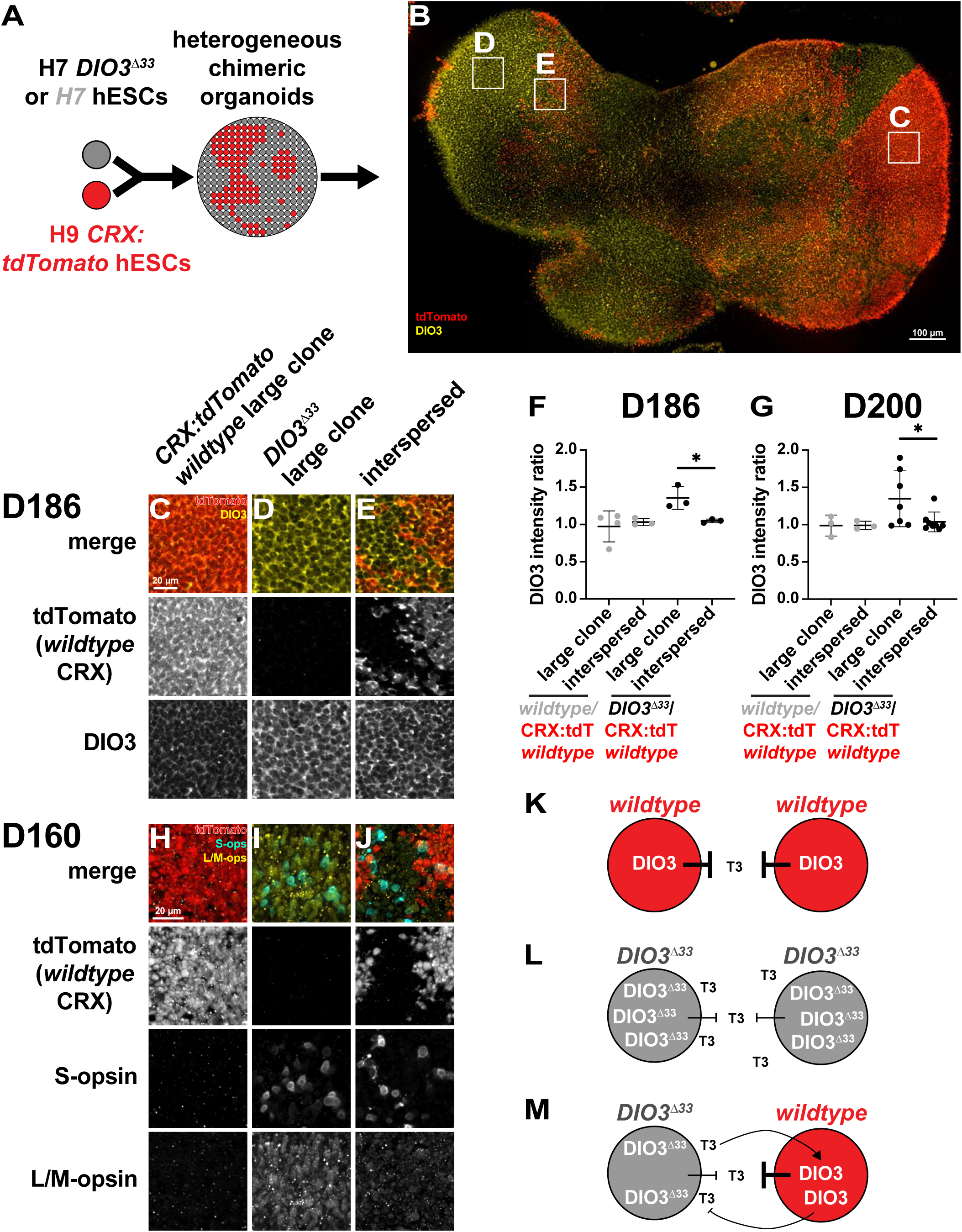
Chimeric organoids reveal cell non-autonomous roles of DIO3. (A) Unlabeled H7 (control) or H7 *DIO3^Δ33^* stem cells were mixed with H9 *CRX:tdTomato* reporter stem cells to generate chimeric organoids. (B) D186 representative chimeric retinal organoid. tdTomato+ regions are H9 *wildtype CRX:tdTomato* cells. tdTomato-regions are H7 *DIO3^Δ33^* cells. Boxed regions correspond with (C-E). (C-E) Clones at D186. (C) large *wildtype CRX:tdTomato* clone, (D) large *DIO3^Δ33^*clone, (E) interspersed cells stained for tdTomato and DIO3. (F) DIO3 intensity ratios for D186 clones. N=3 *wildtype*/*wildtype CRX:tdTomato* chimeric organoids, N=4 *DIO3^Δ33^*/*wildtype CRX:tdTomato* chimeric organoids. N=4 large *wildtype*/*wildtype CRX:tdTomato* matched large clones, N=3 *wildtype*/*wildtype CRX:tdTomato* matched interspersed regions, N=3 *DIO3^Δ33^/wildtype CRX:tdTomato* matched large clones, N=3 *DIO3^Δ33^*:*wildtype* matched interspersed regions. n=50 cells per genotype per clone. Error bars represent SD. Statistics determined by unpaired t test. Unlabeled comparisons are not significant (p>0.05). *DIO3^Δ33^*/*wildtype CRX:tdTomato* large clone vs interspersed ratio p=0.0267. (G) DIO3 intensity ratios for D200 clones. N=5 *wildtype*/*wildtype CRX:tdTomato* chimeric organoids, N=11 *DIO3^Δ33^*/*wildtype CRX:tdTomato* chimeric organoids. N=3 large *wildtype*/*wildtype CRX:tdTomato* matched large clones, N=3 *wildtype*/*wildtype CRX:tdTomato* matched interspersed regions, N=7 *DIO3^Δ33^*/*wildtype CRX:tdTomato* matched large clones, N=8 *DIO3^Δ33^*/*wildtype CRX:tdTomato* matched interspersed regions. n=50 cells per genotype per clone. Error bars represent SD. Statistics determined by unpaired t test. Unlabeled comparisons are not significant (p>0.05). *DIO3^Δ33^*/*wildtype CRX:tdTomato* large clone vs interspersed ratio p=0.0466. (H-J) Representative D160 chimeric organoid stained for S-opsin, L/M-opsin, and tdTomato reporter expression for N=2 organoids which contained all three clone types. (H), large *wildtype CRX:tdTomato* region, (I), large *DIO3^Δ33^*region, (J), boundary/interspersed region. (K-M) Model of cell non-autonomous signaling in (K) homogenous or large *wildtype CRX:tdTomato* regions, (L) homogeneous or large *DIO3^Δ33^* regions, and (M) interspersed regions.

In organoid tissue derived from the *wildtype* cells or *DIO3^Δ33^*mutant cells, cells were unmarked. In organoid tissue derived from the *CRX:tdTomato wildtype* cells, *wildtype* PR precursors and PRs were marked by expression of the *CRX:tdTomato* reporter (Phillips et al., 2018) (**Fig. 5A-B**). This strategy produced chimeric organoids with clones of varying size and location, enabling characterization of spatial effects based on differences in TH signaling regulation in *wildtype* and *DIO3^Δ33^* mutant cells (**Fig. 5B-E**). We observed large clones of >1000 cells of a genotype (**Fig. 5C-D**), and interspersed cells, in which each cell of a genotype was bordered by >1 cell of the other genotype (**Fig. 5E**).

At day 200, DIO3 expression was low in homogenous *wildtype* organoids and higher in homogenous *DIO3^Δ33^* mutant organoids (**Fig. 4E-F**). Within a single organoid, large *wildtype* clones had low DIO3 expression levels, while large *DIO3^Δ33^* clones had higher DIO3 levels (**Fig. 5C-D**), consistent with observations in homogenous organoids of either genotype (**Fig. 4E-F**) and suggesting that any cell non-autonomous effects are distance-limited. In the interspersed cell population, both *wildtype* and *DIO3^Δ33^* mutant cells had similar, intermediate levels of DIO3 (**Fig. 5E**). This result suggests that non-autonomous regulation between cells coupled with feedback results in similar levels of DIO3 protein.

To quantify these differences, we measured DIO3 expression intensity in 50 cells of each genotype per clone type at days 186 and day 200. To control for variability among organoids, we related expression levels with four ratios (**Fig. 5F-G**): (1) *wildtype/CRX:tdTomato wildtype* in large clones, (2) *wildtype/CRX:tdTomato wildtype* in interspersed clones, (3) *DIO3^Δ33^/CRX:tdTomato wildtype* in large clones, and (4) *DIO3^Δ33^/CRX:tdTomato wildtype* in interspersed clones. Ratios (1) and (2) were controls to address differences in *wildtype* DIO3 expression between stem cell lines and were expected to be ∼1. Ratio (3) was a positive control and higher DIO3 expression in the *DIO3^Δ33^* mutant was expected (i.e. >1). Ratio (4), in combination with our controls, allowed for identification of non-cell autonomous signaling effects. A ratio of ∼1 would indicate non-cell autonomous signaling, whereas a ratio >1 would suggest cell autonomous signaling. This approach enabled us to compare ratios of DIO3 expression intensity between organoids (**Fig. 5C-G**). For these experiments, we observed similar results on days 186 and 200 (**Fig. 5F-G**).

In control *wildtype/wildtype CRX:tdTomato* organoids, the ratios of DIO3 expression level intensity in both large and interspersed clones were ∼1 (**Fig. 5F-G**), suggesting that the levels of DIO3 were similar in both clone types and that the H7 and H9 hESC lines had similar DIO3 expression patterns. In contrast, large clones in the *DIO3^Δ33^*/*wildtype CRX:tdTomato* organoids had a greater DIO3 intensity ratio of ∼1.3 (**Fig. 5F-G**), consistent with high DIO3 expression in *DIO3^Δ33^* mutant cells and lower DIO3 expression in *wildtype CRX:tdTomato* cells. Interspersed cells in the *DIO3^Δ33^*/*wildtype CRX:tdTomato* organoids had a DIO3 intensity ratio closer to ∼1 (**Fig. 5F-G**), consistent with non-autonomous mechanisms yielding similar levels of DIO3.

We interpret these data as follows: In homogenous *wildtype* organoids or large *wildtype* clones, functional DIO3 degrades T3, and DIO3 levels are low (**Fig. 5K**). In homogenous *DIO3^Δ33^* mutant organoids or large clones of *DIO3^Δ33^*, DIO3^Δ33^ protein cannot properly degrade T3, and the resulting high T3 environment induces DIO3 expression (**Fig. 5L**). Our observations of intermediate DIO3 levels in interspersed clones of *DIO3^Δ33^*and *wildtype* cells suggest local intercellular feedback regulation. In interspersed *DIO3^Δ33^* cells, impaired DIO3^Δ33^ protein yields high T3, which upregulates DIO3 in neighboring *wildtype* cells. Functional DIO3 in *wildtype* cells degrades T3 to lower T3 levels, leading to downregulation of DIO3 in the *DIO3^Δ33^* cells (**Fig. 5M**). This intercellular feedback coordinates DIO3 expression across cells to yield intermediate levels.

### Cone developmental timing is controlled by cell non-autonomous regulation

To determine if non-autonomous regulation affected PR development, we compared cone subtype development in large *wildtype CRX:tdTomato* clones, large *DIO3^Δ33^* mutant clones, and interspersed cells on day 160 (**Fig. 5H-J, Fig. S6**). Large *wildtype* clones had low photoreceptor density (**Fig. 5H, Fig. S6A, solid boxed region**). Large *DIO3^Δ33^* clones had greater S-opsin+ and L/M-opsin+ cell densities (**Fig. 5I, Fig. S6B, solid boxed region**), consistent with accelerated cone development in *DIO3^Δ33^* cells. Interspersed clones or clone boundaries had an intermediate amount of S-opsin+ and L/M-opsin+ cells (**Fig. 5J, Fig. S6C, solid boxed region**). These data suggest that DIO3 expression and cone subtype developmental timing is regulated by TH levels and locally coordinated across cells.

### TH-mediated regulation of PR specification dictates developmental timing and fate

Our data suggest that the coupling of T3 degradation to cell differentiation—RPCs express DIO3 whereas differentiated neurons do not—implements a feedback loop that controls timing of photoreceptor development. In this “signaling model”, cell ratios and intercellular signaling dynamically tune the rates of cell differentiation. This description contrasts with the probabilistic, cell-intrinsic mechanism previously proposed to control neuronal subtype specification in the retina (“intrinsic model”) (Cepko, 2014).

To identify potential functions of DIO3-mediated feedback, we formalized our observations into a stochastic model of photoreceptor development. In this model (**Fig. 6A**), RPCs differentiate into immature S or L/M cones with T3-dependent rates or differentiate into non-cone fates with a constant rate. Immature cones mature into terminally differentiated cones with constant, T3-independent rates. Finally, S&L/M co-expressing cones arise from immature S or L/M cones via a ‘secondary firing’ of the T3-dependent differentiation program before terminal maturation. T3 levels are regulated by the abundance of undifferentiated RPCs; each RPC expresses DIO3 and is assumed to degrade T3 with first-order kinetics. A detailed discussion of model rate functions, parameters and assumptions can be found in the Supplementary Information.

**Fig. 6.**
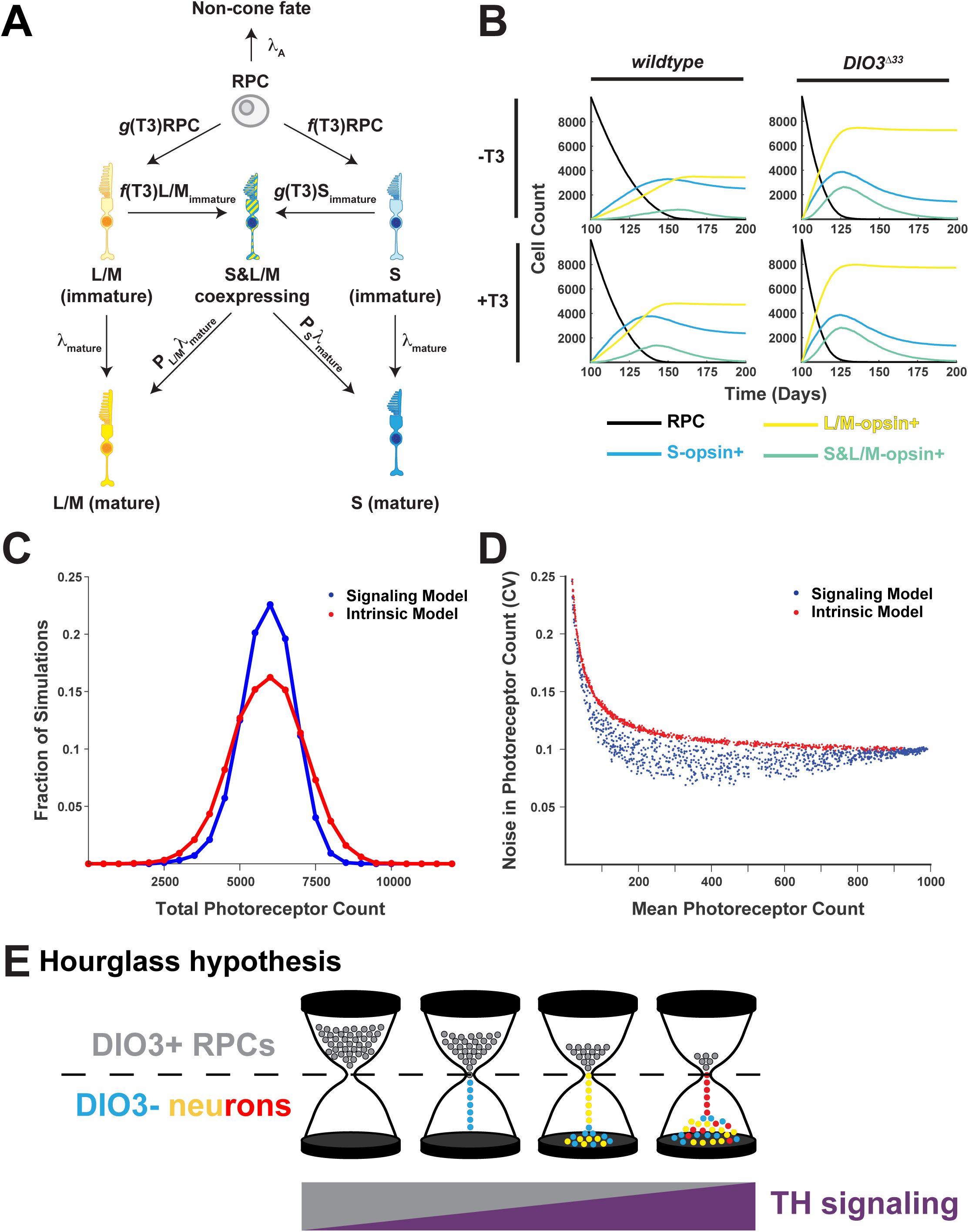
TH feedback confers robustness to photoreceptor specification. (A) Signaling model schematic. RPCs (gray) differentiate into immature L/M (yellow) and S (blue) cones with T3-dependent rates, and non-cone fates with a constant rate. Functions *g* and *f* are selected as Hill functions with n = 1. Immature cones differentiate into mature cones with constant rates. Immature cones can express a second opsin gene to become S&L/M expressing cones (hashed yellow/blue). (B) Simulation of cone specification dynamics. *wildtype* (left) and *DIO3^Δ33^* (right) organoids in the absence (top) and presence (bottom) of exogenous T3 are shown. Curves depict counts of RPCs (black), L/M cones (yellow), S cones (blue), and S&L/M co-expressing cones (green). (C) T3 signaling filters noise in initial RPC abundance. Histograms show the total number of photoreceptors (S + L/M cones) specified in 5000 simulations seeded with random number of initial RPCs in signaling (blue) and cell intrinsic (red) models. (D) Exploration of model parameter space. Simulations (n = 1000) were performed for signaling (blue) and cell-intrinsic (red) models with randomly-sampled parameter values. Each dot represents the performance of 50 replicate simulations with a given parameter set. (E) ‘Hourglass hypothesis’.

Our model captures the key features of photoreceptor specification dynamics in *wildtype* organoids. Specifically, the model reproduces the ordering (S before L/M) of photoreceptor maturation, as well as the appearance of a small number of S&L/M-opsin co-expressing cells (**Fig. 2F-G, 6B**, upper left). To simulate *DIO3^Δ33^* organoid development, we altered the rate constant of DIO3-mediated T3 degradation to 6% of its wild-type value. This modified model correctly predicts the increase in L/M:S cone ratio and accelerated cone specification dynamics observed in *DIO3^Δ33^* organoids (**Fig. 2H-I, 6B,** upper right). Finally, the model recapitulates the observed phenotypes of *wildtype* +T3 (modest acceleration of differentiation) and *DIO3^Δ33^* +T3 (no effect compared to *DIO3^Δ33^*) (**Fig. 2J-M, 6B,** lower left and lower right).

### The TH-dependent mechanism confers developmental robustness

We next used our model to explore whether T3 feedback offers advantages over a cell-intrinsic mechanism. Specifically, we tested whether this control enhances the robustness of photoreceptor specification. To this end, we performed a series of stochastic simulations in which each simulation is seeded with a random number of initial RPCs. We find that T3 signaling enables the system to ‘filter out’ noise in RPC number (**Fig. 6C**); the distribution of final photoreceptors specified in the signaling model (blue) is narrower than that of a cell-intrinsic model (red) in which RPCs stochastically differentiate with constant rates.

Finally, to verify that this noise filtering is a general property of the signaling model—rather than product of specific parameter value choices—we simulated 1000 instantiations of the signaling and intrinsic models, each with randomly-sampled parameter values (**Fig. 6D**). We find that the noise in final photoreceptor count (as measured by the coefficient of variation) is systematically lower for the signaling molecule (blue) than for the intrinsic model (red). Indeed, the cell intrinsic model represents an upper bound for the poorest-performing versions of the signaling model.

## Discussion

The mechanisms that control the timing of human retinal cell development have remained elusive. Our results suggest a link between the temporal regulation of TH signaling and the generation, maturation, and stability of PR subtypes. RPCs express DIO3, which diminishes as PRs and other retinal cells differentiate. As the RPC:neuron ratio decreases, TH degradation is relieved, leading to increased signaling and development of PR subtypes at distinct times. DIO3 and TH signaling levels are coordinated across cells by local intercellular feedback, providing robustness to cone subtype specification compared to probabilistic, intrinsic mechanisms. We discuss the implications of these findings for the specification of PR subtypes and more generally for the strategies that coordinate cell type generation and development in the retina and other contexts.

### DIO3 and TH signaling regulate PR number, developmental timing, and fate stability

The dynamic expression and function of DIO3 are essential for regulating PR development. DIO3 is highly expressed in RPCs and decreases as these cells differentiate into neurons and mature. During retinal development, the asynchronous differentiation of RPCs leads to a progressive reduction in overall DIO3 levels. This reduction establishes a temporal gradient of TH signaling, which provides critical timing cues for PR development.

This temporal gradient mechanism can be regarded as a twist on the “French flag” model of spatial embryonic development, first proposed by Wolpert in 1969 (Wolpert, 1969). In this model, specification is determined by position along a spatial gradient of a morphogen, later shown to be the transcription factor Bicoid in the fly embryo (Driever and Nusslein-Volhard, 1988). In the retina, the gradual accumulation of T3 ‘patterns’ PR development in time, rather than in space.

DIO3 is required to limit PR generation, set PR developmental timing, and ensure PR fate stability. In *DIO3^Δ33^*mutants, S-opsin, L/M-opsin, and Rho are expressed earlier and in greater numbers of cells. Later, subsets of S-opsin+ and Rho+ cells also express L/M opsin, indicating a loss of cell fate exclusivity and a possible transformation in PR subtype identity. These phenotypes are observed in *DIO3^Δ33^*, *wildtype +T3,* and *DIO3^Δ33^ +T3* organoids. The milder phenotype in *wildtype +T3* organoids is consistent with the presence of wildtype DIO3, which is upregulated in high T3 to increase degradation and maintain T3 levels. The highly similar phenotypes of *DIO3^Δ33^* mutant organoids in the presence or absence of additional T3 suggest that DIO3 and TH signaling act in the same pathway, and that T3 levels in the media are saturating.

While a great focus has been placed upon terminal cell fates, the temporality of cell fate decisions is less understood. *DIO3^Δ33^* mutant and *wildtype*+T3 organoids showed an advancement in S-opsin, L/M-opsin, and rhodopsin expression (**Fig. 2, Fig. S2**). This suggests that excess TH signaling accelerates cell fate specification. This could be due to either an advancement of cell cycle exit or a preference towards differentiating photoreceptors from the RPC pool. TH has previously been shown to influence cell cycle (Durand and Raff, 2000). In oligodendrocyte precursor cells, TH signaling promotes differentiation and is hypothesized to be part of a timing mechanism in which cells count how many cell cycles occur before terminal differentiation (Alisi et al., 2004; Durand and Raff, 2000; Porlan et al., 2004). If a similar mechanism is occurring in PR precursors, the excess TH in *wildtype*+T3 and in *DIO3^Δ33^* might be inducing cell cycle exit prematurely, causing the advancement of opsin+ cells.

The co-expression of L/M-opsin in S-opsin+ cells in *DIO3^Δ33^*mutant and *wildtype* +T3 organoids mirrors our previous observations in the developing foveola (Hussey et al., 2024), the central region of the retina responsible for high-acuity vision. The presumptive foveola contains S-opsin+ cells during early fetal development, co-expression of L/M-opsin in S-opsin+ cells later, and L/M cones in adults (Cornish et al., 2004a; Hussey et al., 2024; Xiao and Hendrickson, 2000). This cell fate progression is associated with the sustained high expression of DIO2, an enzyme that generates T3 and promotes TH signaling. The co-expression of L/M-opsin in S-opsin+ cells, followed by the loss of S-opsin+ cells (Hussey et al., 2024), is consistent with a transformation in cell fate.

In addition to cone development, rod development is also regulated by DIO3 and TH signaling, which had not been observed previously. Rod development has been previously shown to be regulated by retinoic acid (RA), another nuclear hormone signaling mechanism. RA promotes rod specification in human retinal organoids, mice, and zebrafish (Kelley et al., 1999; Khanna et al., 2006; Oh et al., 2008) (Prabhudesai et al., 2005). Our data suggest that TH works alongside RA to control the timing of rod development. RA receptors and TH receptors can heterodimerize (Butler and Parker, 1995; Koenig, 1998; Roberts et al., 2006; Zhang et al., 1992) and we hypothesize that modulation of both RA and TH signaling are important for developmental timing and subtype specification of PRs.

We previously showed that RA promotes early human cone subtype fates, specifying S cone fate over L/M cone fate and M cone fate over L cone fate (Hadyniak et al., 2024; Hussey et al., 2024). Consistent with these roles for RA signaling, aldehyde dehydrogenases (ALDHs), enzymes that generate RA, are highly expressed early in human retinal development (Hadyniak et al., 2024; Hoshino et al., 2017). Both RA-generating ALDHs and TH-degrading DIO3 are highly expressed early and decrease during development suggesting high RA signaling early transitions to high TH signaling late, leading to temporally inverse gradients of signaling. These temporal counter gradients are conceptually similar to the spatial counter gradients of Shh and BMP/Wnt that pattern the neurons of the spinal cord (Briscoe et al., 2001; Briscoe and Ericson, 1999; Briscoe et al., 2000; Dessaud et al., 2010; Douceau et al., 2023; Ericson et al., 1997a; Ericson et al., 1997b; Exelby et al., 2021; Fuccillo et al., 2006; Jessell, 2000; Patten and Placzek, 2002; Stamataki et al., 2005). Since both TH and RA act through a diverse set of nuclear hormone receptors to activate and repress gene expression (Aranda and Pascual, 2001; Astapova and Hollenberg, 2013; Cho et al., 2023), these counter-gradients may work together to trigger temporal events during retinal development.

Beyond the retina, the temporal regulation of TH signaling plays important roles in the development of several other tissues. During development of the cochlea in mouse, TH is limited early and increases before the onset of hearing (Campos-Barros, 1999; Ng et al., 2004; Ng et al., 2009). In the brain, a similar transition from low to high TH signaling regulates differentiation of oligodendrocytes (Ahlgren et al., 1997; Barres et al., 1994; Billon et al., 2001; Calza et al., 2002; Carre et al., 1998; Courtin et al., 2005; Gao et al., 1998; Guadano-Ferraz et al., 1997; Lee and Petratos, 2016; Rodriguez-Pena, 1999; Tu et al., 1997; Tu et al., 1999). Our findings in the retina may be a paradigm for understanding the mechanisms controlling the temporality of TH signaling and development in other contexts.

### Comparing roles for DIO3 in human and mouse PR development

Our mechanistic studies of DIO3 reveal similarities and differences in its role in PR development in human retinal organoids and mouse retinas. In mouse retinas, human retinas, and human retinal organoids, *DIO3* is highly expressed early and gradually decreases over time (**Fig. 1**) (Eldred et al., 2018; Ng et al., 2017; Ng et al., 2010). *Dio3Δ* mutant mice display cone loss (Lu et al., 2009; Ma et al., 2014; Ng et al., 2009; Ng et al., 2017; Ng et al., 2011; Ng et al., 2010; Roberts et al., 2006). Similarly, *DIO3* null mutant organoids fail to differentiate retinal tissue (**Fig. S1E-H**). In contrast, *DIO3^Δ33^*mutant organoids differentiate retinal tissue with accelerated PR development (**Fig 2, 3**).

These differences partly stem from the *in vivo* vs. *in vitro* conditions. In mice, TH signaling is influenced by circulating TH levels and interactions with neighboring tissues. In contrast, organoids are exposed to a consistent TH supply provided by the media, meaning any changes in TH signaling must originate from retinal organoid tissue-derived mechanisms.

Additionally, the nature of the mutant alleles contributes to the phenotypic differences. Null mutants lead to loss of retinal cells or failure to differentiate retinal tissue. In contrast, *DIO3^Δ33^* mutant organoids which exhibit a significant yet incomplete loss of function, successfully differentiate retinal tissue, revealing new roles for DIO3-mediated regulation of PR development.

### Differentiation-dependent control of neuronal subtype developmental timing

Our data suggest that the ratio of progenitors and neurons controls PR developmental timing. As this ratio decreases, the proportion of cells expressing DIO3 decreases, leading to increased TH signaling. This mechanism shares some similarities with the regulation of retinal ganglion cell (RGC) generation. RGCs are the first neurons to develop in the retina. Studies in mice have shown that RCGs express Sonic Hedgehog (Shh), which signals to RPCs to inhibit RGC production (Stenkamp and Frey, 2003; Wang et al., 2005; Zhang and Yang, 2001). This mechanism provides a shutoff switch for RGC production. As the population of Shh-expressing RGCs increases, RPC differentiation into RGCs is stopped, allowing the differentiation of other cell types to progress.

Similar to the relationship between Shh signaling and RGC development, our study identified a signaling regulatory mechanism linked to progenitor differentiation. For RGCs, differentiation leads to increased Shh signaling, which in turn inhibits further RGC differentiation. For PRs, differentiation leads to reduced DIO3 expression and activity, increasing TH signaling to promote PR development and terminal L/M cone fate.

Other tissues also use differentiation-regulated signaling to temporally drive fate choices. In the spinal cord, early-born medial lateral motor column (LMC) neurons express RALDH2, which produces RA, which induces late-born LMC to adopt a lateral LMC fate (Sockanathan and Jessell, 1998). Thus, early-born neurons express a signal that specifies late-born neurons. In contrast, in the retina, the generation of neurons relieves degradation of a signal that drives PR development.

### Signaling and feedback promote robustness of photoreceptor developmental timing

TH signaling is regulated by negative feedback to maintain homeostatic levels on the organismal scale. The HPT axis responds to extremes in TH levels by modulating TSH and TRH release from the hypothalamus and pituitary, respectively, to control TH synthesis and release. Here, we present evidence for local intercellular feedback and coordination of TH regulation in retinal development. The decreasing ratio of progenitors and neurons controls the levels of TH signaling to regulate the temporality of PR development. Signaling is coordinated by feedback between and within retinal cells, which are sensitive to TH levels and respond by modulating expression of TH regulators.

Our experimental results and modeling suggest that coupling TH signaling to photoreceptor differentiation may confer robustness to retinal development (**Fig. 6C-D**). The signaling model achieves tight timing of the duration of photoreceptor specification (**Fig. 6B-D**)— a potentially useful property for coordinating the development of two independent eyes. As we explore in the supplementary information, the principles underlying this noise control are also employed to achieve precise control in biofilm development (Lord et al., 2019; Norman et al., 2013) and synthetic oscillators (Potvin-Trottier et al., 2016).

Whereas our findings identify new roles for signaling in the temporal regulation of retinal PR development, these extrinsic mechanisms must work together with intrinsic mechanisms involving chromatin and transcriptional regulation to differentiate retinal cell types (Andzelm et al., 2015; Cepko, 2014; Emerson et al., 2013; La Torre et al., 2013; Schick et al., 2019; Wang et al., 2014).

### The hourglass hypothesis for regulation of photoreceptor developmental timing

Our data suggest that the changing ratio of RPCs to neurons regulates DIO3 levels, which in turn control T3 levels to determine the timing of PR development. To conceptualize this mechanism, we propose an ‘hourglass hypothesis’ (**Fig. 6E**). In this analogy, the sand at the top of the hourglass represents progenitors, while the sand at the bottom represents differentiated retinal cells. Initially, the hourglass is full at the top, meaning the retina is composed primarily of progenitors. These progenitors express DIO3, maintaining low TH signaling. As development progresses, the sand in the hourglass trickles downward as the progenitors gradually differentiate. This shift reduces the progenitor-to-differentiated cell ratio, decreasing the proportion of cells that express DIO3 and increasing TH signaling. As TH levels reach critical thresholds, PR subtypes are generated and develop at specific times. Eventually, all the sand reaches the bottom, signifying the depletion of the progenitor pool and the complete generation of all retinal neurons and Müller glia.

### Implications for retinal therapies

Our work has identified how cell-population-based signaling coordinates developmental timing in the retina. These studies not only deepen our understanding of cell fate specification and human retinal development but also holds potential implications for translational medicine.

Globally, thyroid disorders affect ∼200 million people worldwide (2012). As TH affects many aspects of development, multiple layers of regulation protect the fetus from fluctuations in maternal TH and fetal TH as the fetal HPT develops. This complexity makes it difficult to interrogate the crucial role of TH in retinal development in a complex organism. A strength of our retinal organoid experiments is the ability to isolate the developing retinal tissue from systemic TH regulation. This enabled us to examine the role of TH regulation in retinal development without complication from the HPT axis. Similarly, human retinal organoids have been used to identify mechanisms of retinal development and model degeneration (Cuevas et al., 2021; Fligor et al., 2018; Guy et al., 2021; Kallman et al., 2020; Kandoi et al., 2024; Lu et al., 2020; O’Hara-Wright and Gonzalez-Cordero, 2020; Phillips et al., 2014; Sridhar et al., 2020).

Technological advances in organoids combined with deeper understanding of developmental processes can advance cell and gene therapies. TH dysregulation has been suggested to have a role in several retinal diseases including diabetic retinopathy and age-related macular degeneration (Nicolini et al., 2024) and further study may reveal new targets for therapeutics. Additionally, retinal organoids can be used as a source of transplantable, healthy retinal cells (Gasparini et al., 2022; Iwama et al., 2024; Li et al., 2021; Shirai et al., 2016; Singh et al., 2019; Watari et al., 2023; Zou et al., 2019). An understanding of mechanisms of cell fate specification can be leveraged to generate organoids with cell constituencies tailored to regions of the retina such as the fovea, which is enriched in L/M cones (Bumsted and Hendrickson, 1999; Cornish et al., 2004a; Cornish et al., 2004b; Diaz-Araya and Provis, 1992; Hussey et al., 2024).

## Acknowledgments

CLM was supported by NIH F31EY033187. KCE was supported by the Damon Runyon Cancer Research Foundation DRG 32-20 and a Hanna H. Gray Fellows Program Award from the Howard Hughes Medical Institute Grant GT15994. IAG is supported by NICHD Grant R24HD000836. CPS was supported by the Visual Sciences Training grant 2T32EY007143. TAR was supported by the Foundation Fighting Blindness TA-RM-0620-0788-463 UWA. SB was supported by the NEI Grants R01EY036173 and R01EY031685. RJJ was supported by National Eye Institute R01EY030872, the BrightFocus foundation G2019300, and the Maryland Technology Development Corporation 2022-MSCRFD-5895.

---

**Supplemental Fig. 1.**
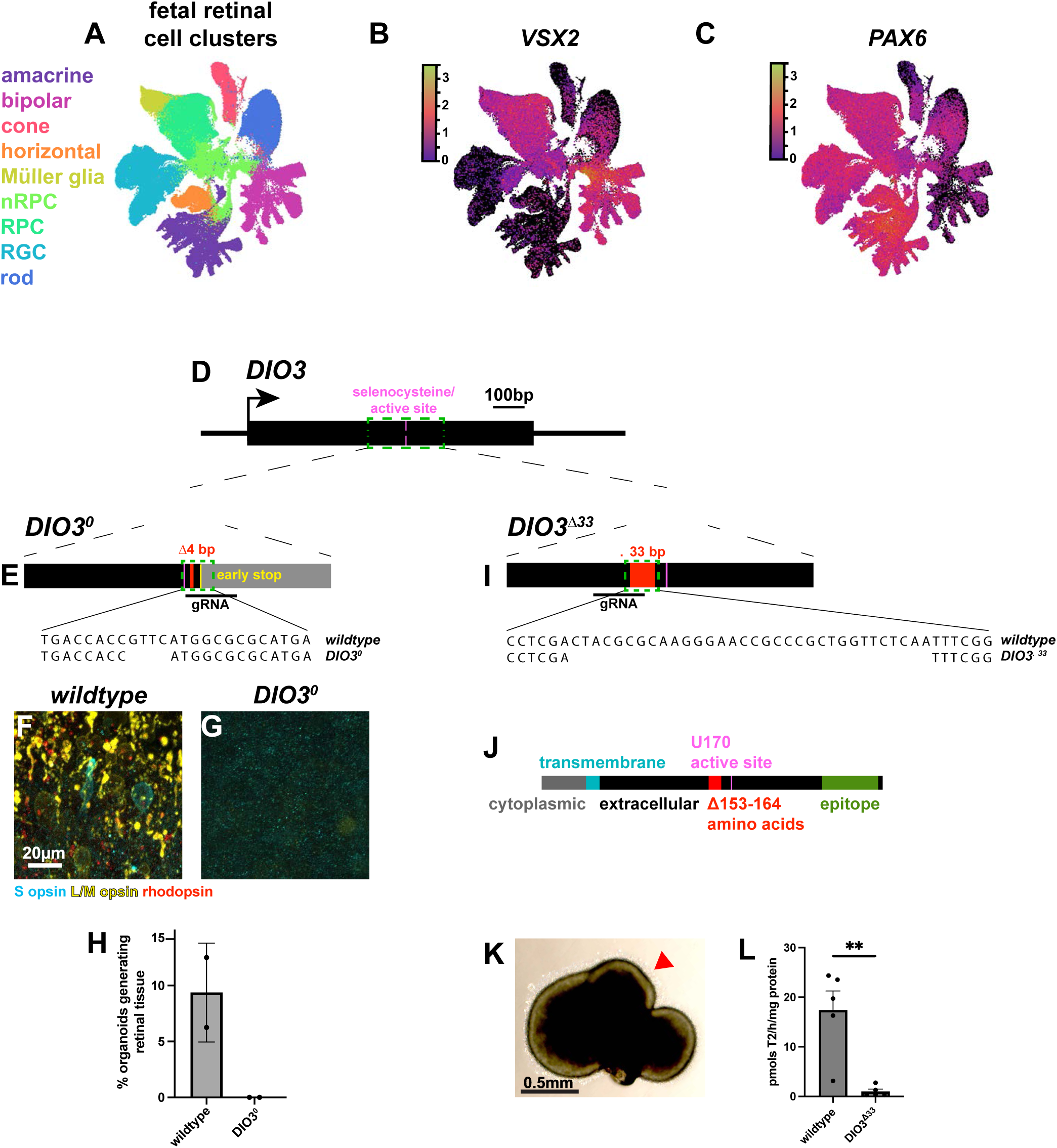
Identification of fetal RPC clusters, generation and characterization of DIO3 mutants. (A) UMAP from CZ CELLxGENE Discover (Zuo et al., 2024) of human fetal scRNA-seq with major cell types labeled, as in Fig. 1A. (B) *VSX2* expression in fetal human retina limited to *VSX2*+ cells, overlaid on the unannotated UMAP. (C) *PAX6* expression in fetal human retina limited to *PAX6*+ cells, overlaid on the unannotated UMAP. (D) *DIO3* locus. Selenocysteine active site marked in pink. Zoom in on green dashed region in **Fig. S1E** and **Fig. S1I**. (E) *DIO3^0^* mutation. The 4 base pair deletion (red) induced an early stop (yellow). Below, the *wildtype* and *DIO3^0^* sequences of this region. (F-G) Day 200 *wildtype* (F) and *DIO3^0^* (G) stained for S-opsin, L/M-opsin, rhodopsin. (H) Percent of *wildtype* and *DIO3^0^* organoids generating retinal tissue. N=384 organoids, 2 independent differentiations, each with *wildtype* and *DIO3^0^* grown in parallel. (I) *DIO3^Δ33^*with a 33 base pair in-frame deletion (red). Below, the wildtype and *DIO3^Δ33^*allele sequences. (J) Protein sequence of wildtype DIO3. Teal = transmembrane region. Magenta = selenocysteine active site. Grey = cytoplasmic domain. Black = extracellular domain. Red = Δ33bp/11amino acid region deleted in *DIO3^Δ33^* mutation. (K) *DIO3^Δ33^* organoid at day 200 at 4x magnification. Red arrowhead indicates photoreceptor outer segments. (L) Deiodination activities in *wildtype* and *DIO3^Δ33^* organoids at day 76. Error bars represent SEM. p value= 0.0029 by unpaired t-test.

**Supplemental Fig. 2.**
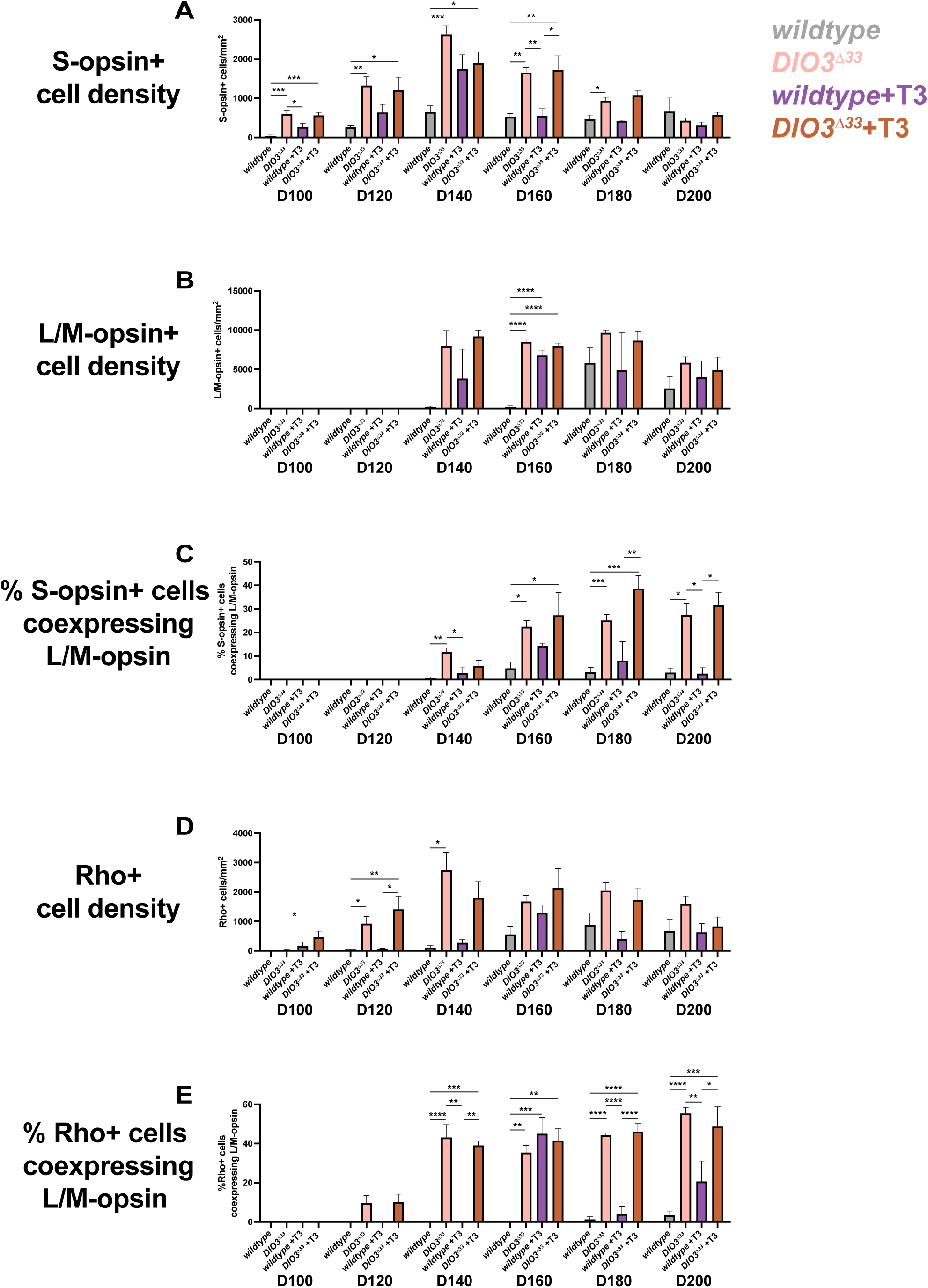
Comparison of photoreceptor development over time. Data as in Fig. 2G, I, K, M, **S3Q-T,** re-plotted to compare per timepoint across conditions. Graph colors consistent with Fig. 2E per condition. Error bars represent SEM, statistics determined via one-way ANOVA with Tukey’s multiple comparisons test unless otherwise noted. Unlabeled comparisons are not significant (p>0.05). (A) S-opsin+ cell density. D100 ANOVA p=0.0002 *wildtype* vs *DIO3^Δ33^* p= 0.0002 *wildtype* vs *DIO3^Δ33^*+T3 p=0.0009 *wildtype* +T3 vs *DIO3^Δ33^* p=0.022 D120 ANOVA p=0.002 *wildtype* vs *DIO3^Δ33^* p=0.0015 *wildtype* vs *DIO3^Δ33^* +T3 p=0.0482 D140 ANOVA p=0.0003 *wildtype* vs *DIO3^Δ33^* p=0.0002 *wildtype* vs *DIO3^Δ33^* +T3 p=0.0334 D160 ANOVA p=0.0004 *wildtype* vs *DIO3^Δ33^* p=0.0024 *wildtype* vs *DIO3^Δ33^* +T3 p=0.0077 *wildtype* +T3 vs *DIO3^Δ33^* p= 0.0072 *wildtype* +T3 vs *DIO3^Δ33^* +T3 p=0.0159 D180 ANOVA p=0.0074 *wildtype* vs *DIO3^Δ33^* p= 0.0197 D200 ANOVA p=0.4456 (ns). (B) L/M-opsin+ cell density. D100 ANOVA p=0.1604 (ns) D120 ANOVA p=0.0556 (ns) D140 ANOVA p=0.0556 (ns) D160 ANOVA p<0.0001 *wildtype* vs *wildtype* +T3 p<0.0001 *wildtype* vs *DIO3^Δ33^* p<0.0001 *wildtype* vs *DIO3^Δ33^* +T3 p<0.0001 D180 ANOVA p=0.0312 (ns) D200 ANOVA p=0.2987 (ns) (C) % S-opsin+ cells co-expressing L/M-opsin. D140 ANOVA p=0.0034 *wildtype* vs *DIO3^Δ33^* p=0.0043 *wildtype* +T3 vs *DIO3^Δ33^* p=0.0415; D160 ANOVA p=0.0251 *wildtype* vs *DIO3^Δ33^* p=0.0419 *wildtype* vs *DIO3^Δ33^* +T3 p=0.0313 D180 ANOVA p<0.0001 *wildtype* vs *DIO3^Δ33^* p= 0.0003 *wildtype* vs *DIO3^Δ33^* +T3 p=0.0001 *wildtype* +T3 vs *DIO3^Δ33^* +T3 p=0.0085 D200 ANOVA p=0.0053 *wildtype* vs *DIO3^Δ33^* p=0.0386 *wildtype* +T3 vs *DIO3^Δ33^* p=0.034 *wildtype* +T3 vs *DIO3^Δ33^* +T3 p=0.0469 (D) Rho+ cell density D100 Kruskal-Wallis with Dunn’s multiple comparisons test p=0.0197 *wildtype* vs *DIO3^Δ33^* +T3 p= 0.0171 D120 ANOVA p=0.0021 *wildtype* vs *DIO3^Δ33^* p=0.0159 *wildtype* vs *DIO3^Δ33^* +T3 p=0.007 *wildtype* +T3 vs *DIO3^Δ33^* +T3 p=0.0307 D140 ANOVA p=0.0165 *wildtype* vs *DIO3^Δ33^* p=0.0259 D160 ANOVA p=0.0573 (ns) D180 ANOVA p=0.0527 (ns) D200 ANOVA p=0.118 (ns) (E) % Rho+ cells co-expressing L/M-opsin. D100 Kruskal-Wallis with Dunn’s multiple comparisons test p>0.9999 D120 Kruskal-Wallis with Dunn’s multiple comparisons test p=0.0334 D140 ANOVA p<0.0001 *wildtype* vs *DIO3^Δ33^* p<0.0001 *wildtype* vs *DIO3^Δ33^* +T3 p=0.0009 *wildtype* +T3 vs *DIO3^Δ33^* p=0.0017 *wildtype* +T3 vs *DIO3^Δ33^* +T3 p=0.0072 D160 ANOVA p=0.0007 *wildtype* vs *wildtype* +T3 p=0.0008 *wildtype* vs *DIO3^Δ33^* p=0.0019 *wildtype* vs *DIO3^Δ33^* +T3 p=0.0017 D180 ANOVA p<0.0001 *wildtype* vs *DIO3^Δ33^* p<0.0001 *wildtype* vs *DIO3^Δ33^*+T3 p<0.0001 *wildtype* +T3 vs *DIO3^Δ33^* p<0.0001 *wildtype*+T3 vs *DIO3^Δ33^* +T3 p<0.0001 D200 ANOVA p<0.0001 *wildtype* vs *DIO3^Δ33^* p<0.0001 *wildtype* vs *DIO3^Δ33^* +T3 p=0.0007 *wildtype* +T3 vs *DIO3^Δ33^* p=0.0023 *wildtype* +T3 vs *DIO3^Δ33^* +T3 p=0.0443

**Supplemental Fig. 3.**
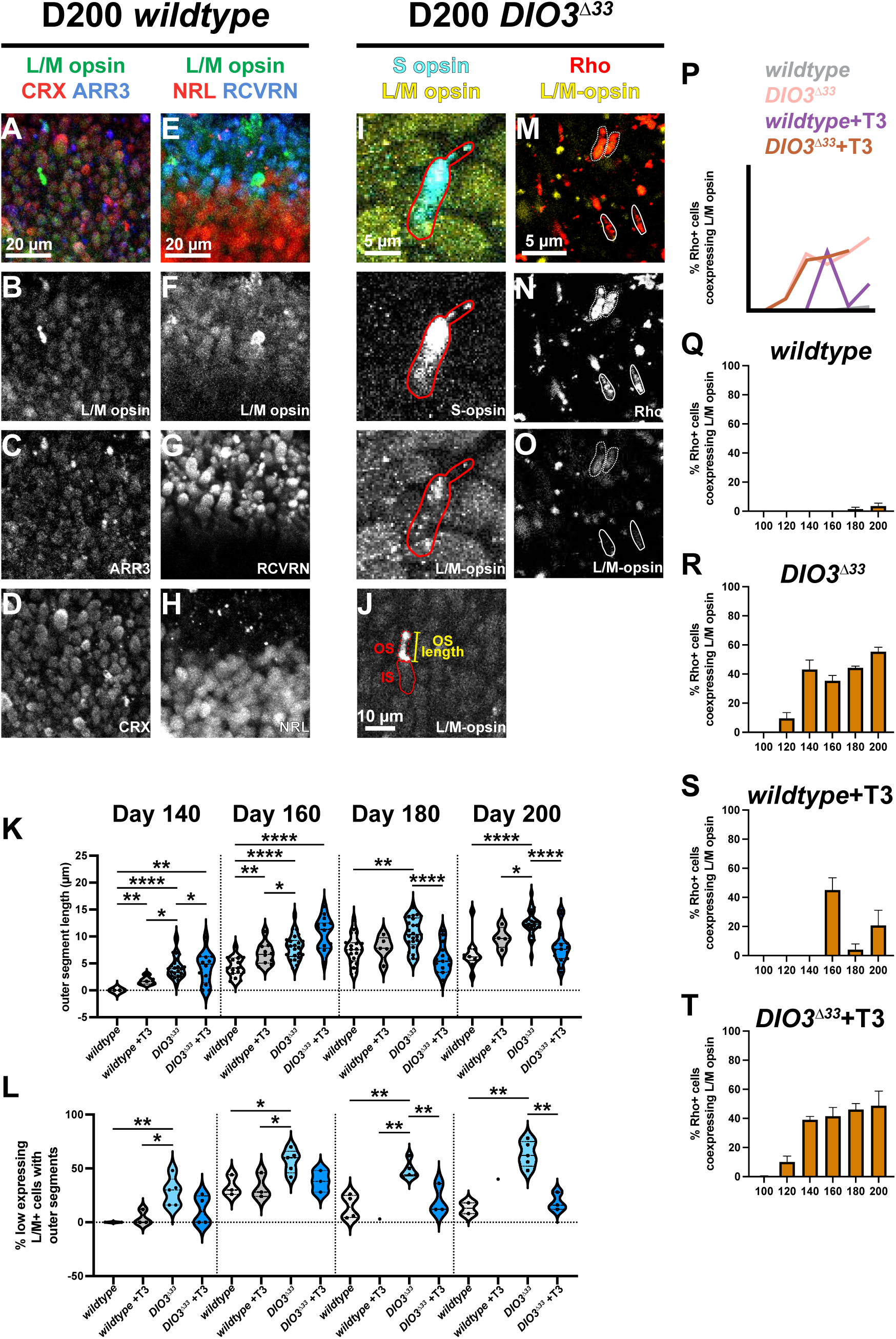
Low L/M-opsin+ cells express cone genes, display cone morphologies, and can co-express other opsins. (A-H) Expression of photoreceptor markers in *wildtype* day 200 organoids. (A-C) Co-expression of low expressing L/M-opsin+ cells with CRX and ARR3. (E-H) Co-expression of L/M-opsin with RCVRN but not NRL. (I) *DIO3^Δ33^* day 200 S-opsin+ cell co-expressing L/M-opsin+ cell. Red outline delineates cell boundary. (J) Low L/M-opsin+ expressing cell in *DIO3^Δ33^* day 200 organoid with an inner segment (IS) and outer segment (OS). OS length (yellow) measured for (K). (K-L) Outer segment length (K) and % of low expressing L/M-opsin+ cells with OS (L) in *wildtype* (light grey), *wildtype*+T3 (dark grey), *DIO3^Δ33^* (light blue), *DIO3^Δ33^* +T3 (dark blue) organoids at day 140, day 160, day 180, and day 200. Error bars represent SEM, two-tailed t-test performed to determine statistical significance. Unlabeled comparisons are not significant (p>0.05). *wildtype* D140 N=4, D160 N=3, D180 N=4, D200 N=2; *wildtype* +T3 D140 N=1, D160 N=3, D180 N=1, D200 N=1. *DIO3^Δ33^* D140 N=5, D160 N=5, D180 N=5, D200 N=5. *DIO3^Δ33^* +T3 D140 N=4, D160 N=3, D180 N=3, D200 N=3. (K) D140: *wildtype* vs *wildtype* +T3 p= 0.002, *wildtype* vs *DIO3^Δ33^* p=0.0006, *wildtype* vs *DIO3^Δ33^* +T3 p=0.0141, *wildtype* +T3 vs *DIO3^Δ33^* p=0.0187. D160: *wildtype* vs *wildtype* +T3 p= 0.0042, *wildtype* vs *DIO3^Δ33^* p<0.0001, *wildtype* vs *DIO3^Δ33^* +T3 p<0.0001, *wildtype* +T3 vs *DIO3^Δ33^* +T3 p=0.0015, *DIO3^Δ33^* vs *DIO3^Δ33^* +T3 p=0.0035. D180: *wildtype* vs *DIO3^Δ33^* p=0.001, *DIO3^Δ33^*vs *DIO3^Δ33^* +T3 p<0.0001. D200: *wildtype* vs *DIO3^Δ33^* p<0.0001, *wildtype* +T3 s *DIO3^Δ33^* p=0.046, *DIO3^Δ33^* vs *DIO3^Δ33^* +T3 p<0.0001. n=5 cells of this morphology analyzed per organoid per timepoint and condition. (L) D140: *wildtype* vs *DIO3^Δ33^*p=0.004, *wildtype*+T3 vs *DIO3^Δ33^* p=0.0278. D160 *wildtype* vs *DIO3^Δ33^* p=0.0218, *wildtype* +T3 vs *DIO3^Δ33^*p=0.0255. D180: *wildtype* vs *DIO3^Δ33^* p=0.0011, *wildtype* +T3 vs *DIO3^Δ33^* p=0.0071, *DIO3^Δ33^* vs *DIO3^Δ33^* +T3 p=0.0097. D200 *wildtype* vs *DIO3^Δ33^* p=0.0031, *DIO3^Δ33^*vs *DIO3^Δ33^* +T3 p=0.0014. (M-O) Single z image of cells that co-express Rho and L/M-opsin in a *DIO3^Δ33^* organoid at day 200. Solid outline marks Rho+ only cells, dashed outline represents Rho&L/M-opsin co-expressing cells. (P) Graphical representation of (P-T) quantifications. (Q-T) Quantification of %Rho+ cells expressing L/M-opsin in (Q) *wildtype*, (R) *wildtype* +T3, (S) *DIO3^Δ33^*, and (T) *DIO3^Δ33^*+T3. Organoids without rods were excluded from the analysis. Error bars represent SEM. *Wildtype* D100 N=0, D120 N=3, D140 N=5, D160 N=3, D180 N=3, D200 N=4. *DIO3^Δ33^* D100 N=2, D120 N=8, D140 N=6, D160 N=10, D180 N=12, D200 N=9. *wildtype* +T3 D100 N=2, D120 N=2, D140 N=2, D160 N=4, D180 N=2, D200 N=3. *DIO3^Δ33^*+T3 D100 N=4, D120 N=3, D140 N=4, D160 N=4, D180 N=3, D200 N=3.

**Supplemental Fig. 4.**
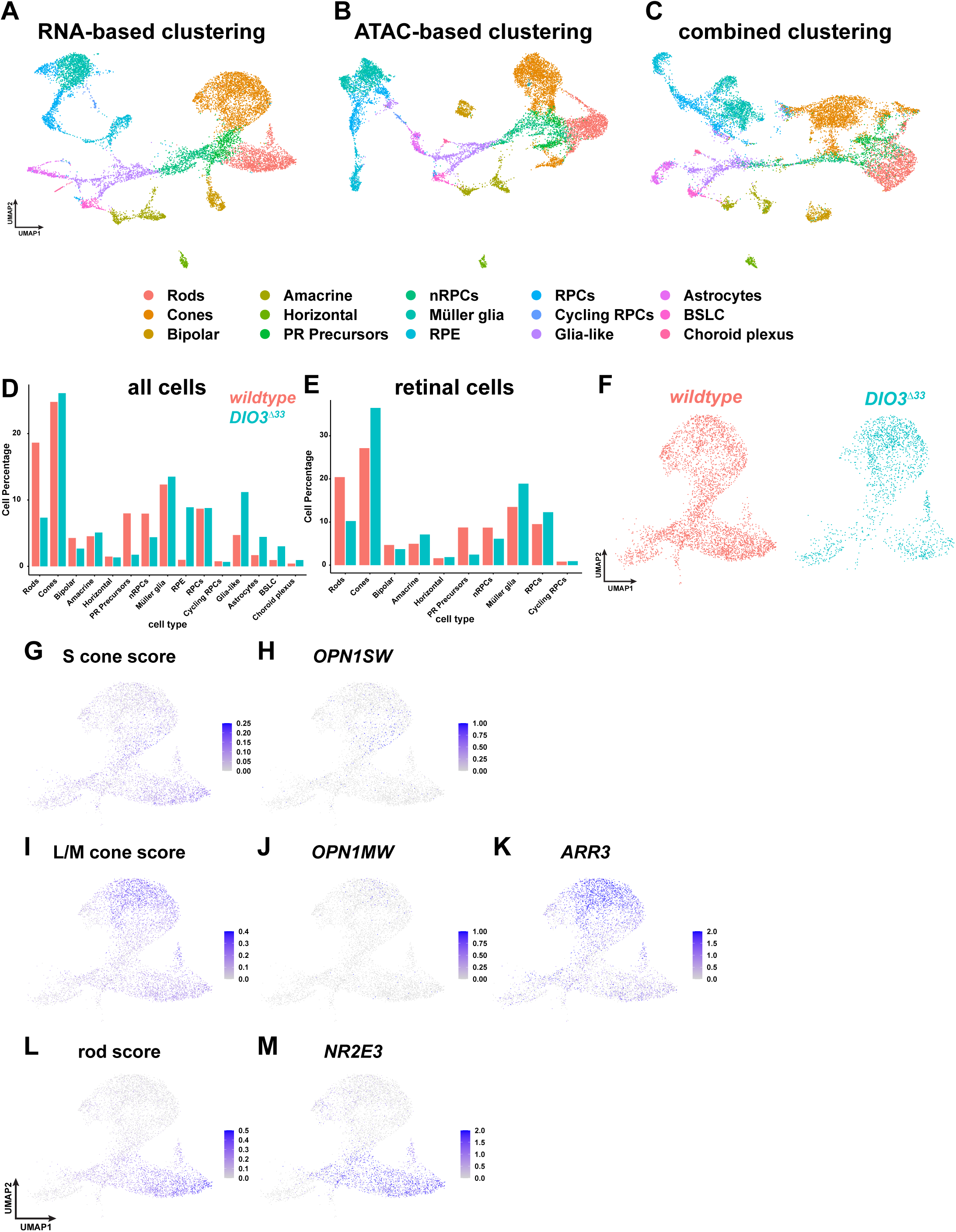
Characterization of *DIO3^Δ33^* mutant organoids with single nucleus multiomics. (A) UMAP clustering of *wildtype* and *DIO3^Δ33^* mutant organoids based on single nucleus RNA-sequencing. (B) UMAP clustering of *wildtype* and *DIO3^Δ33^* mutant organoids based on single nucleus ATAC-sequencing. (C) UMAP clustering of *wildtype* and *DIO3^Δ33^* mutant organoids with contributions from both the RNA- and ATAC-sequencing. (D) Cell proportions for all cells in *wildtype* (coral) and *DIO3^Δ33^* (teal) organoids. (E) Cell proportions for all retinal cells in *wildtype* (coral) and *DIO3^Δ33^* (teal) organoids. (F) UMAP of photoreceptor lineage cells (nRPCs, PR precursors, cones, rods) in *wildtype* (coral) and *DIO3^Δ33^* (teal) organoids. (G-M) Gene expression heatmaps overlaid on (F) UMAP. (G) Compiled S-cone score, (H) *OPN1SW*, (I) L/M-cone score, (J) *OPN1MW*, (K) *ARR3* (L) rod score, and (M) *NR2E3*.

**Supplemental Fig. 5.**
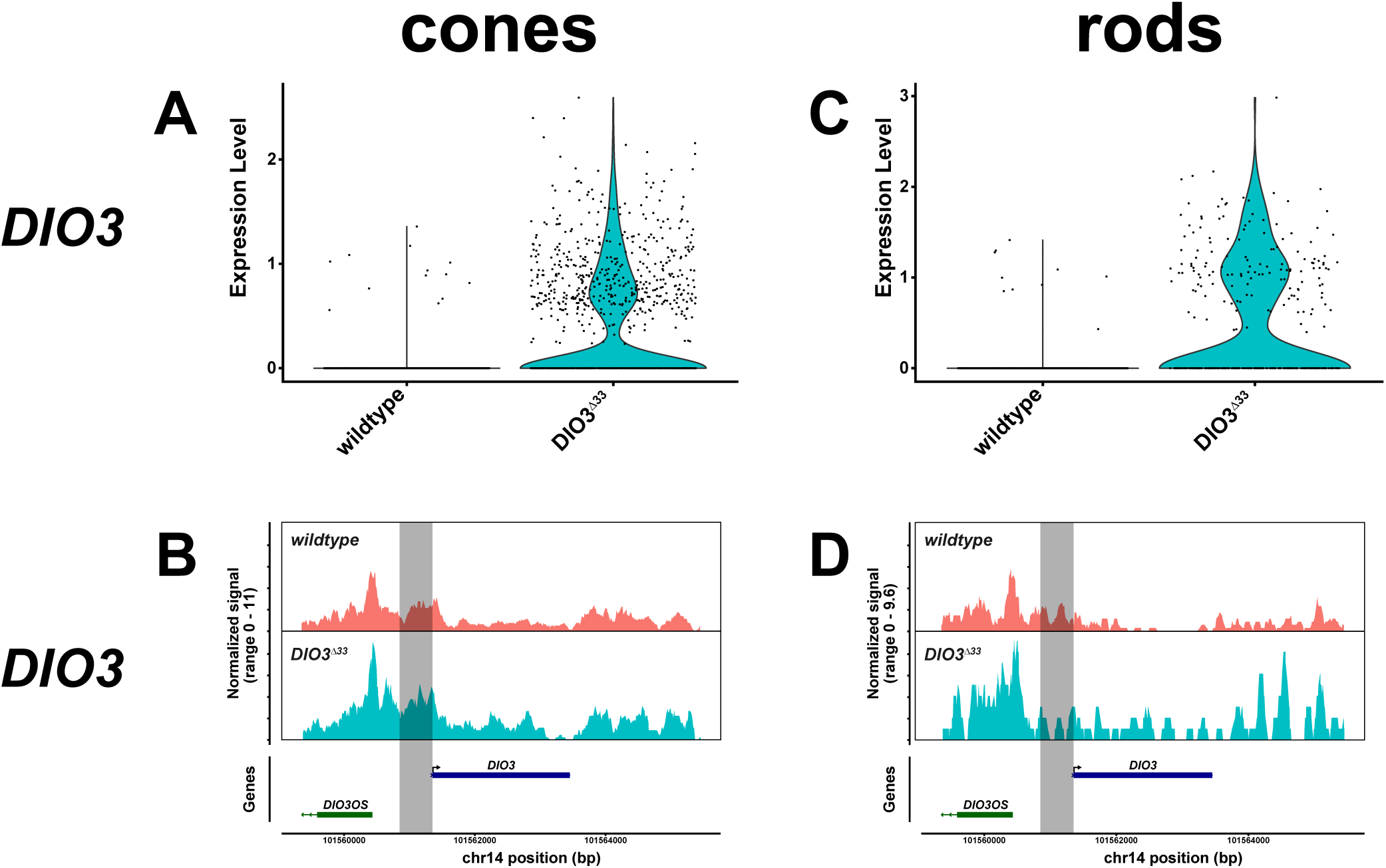
Accessibility and expression of TH regulatory genes in photoreceptors. (A, C) Expression of *DIO3* in cones (A) and rods (C) in *wildtype* and *DIO3^Δ33^* organoids. (B, D) Accessibility of *DIO3* in cones (B) and rods (D) in *wildtype* and *DIO3^Δ33^* organoids.

**Supplemental Fig. 6.**
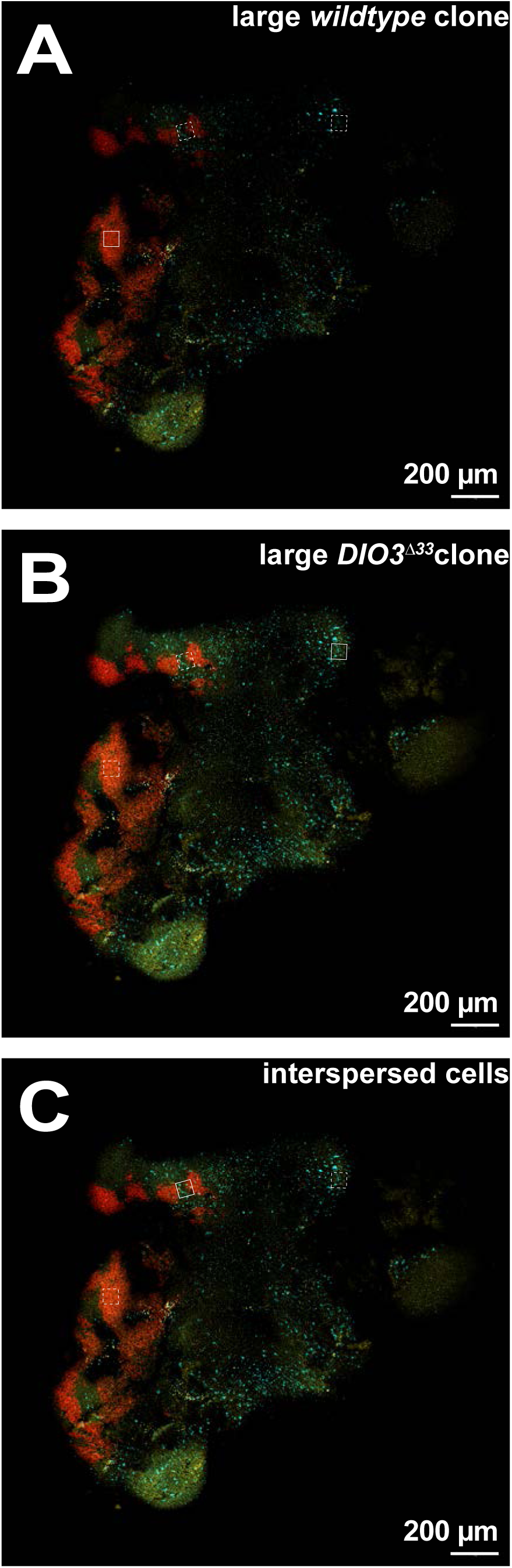
Methods of organoid differentiation. (A-C) D160 chimeric retinal organoid as in Fig 5 H-J. tdTomato+ regions are H9 *wildtype CRX:tdTomato* cells. tdTomato+ regions are H7 *DIO3^Δ33^* cells. Due to the 3D shape of the organoid, images were taken from three different focal planes. Solid box indicates region of interest in that focal plane. Dotted box indicates region of interest from a different focal plane. Solid boxed region in A correspond with **Fig 5H**. Boxed region in B correspond with **Fig 5I**. Boxed region in C correspond with **Fig 5J**.

**Supplemental Fig. 7.**
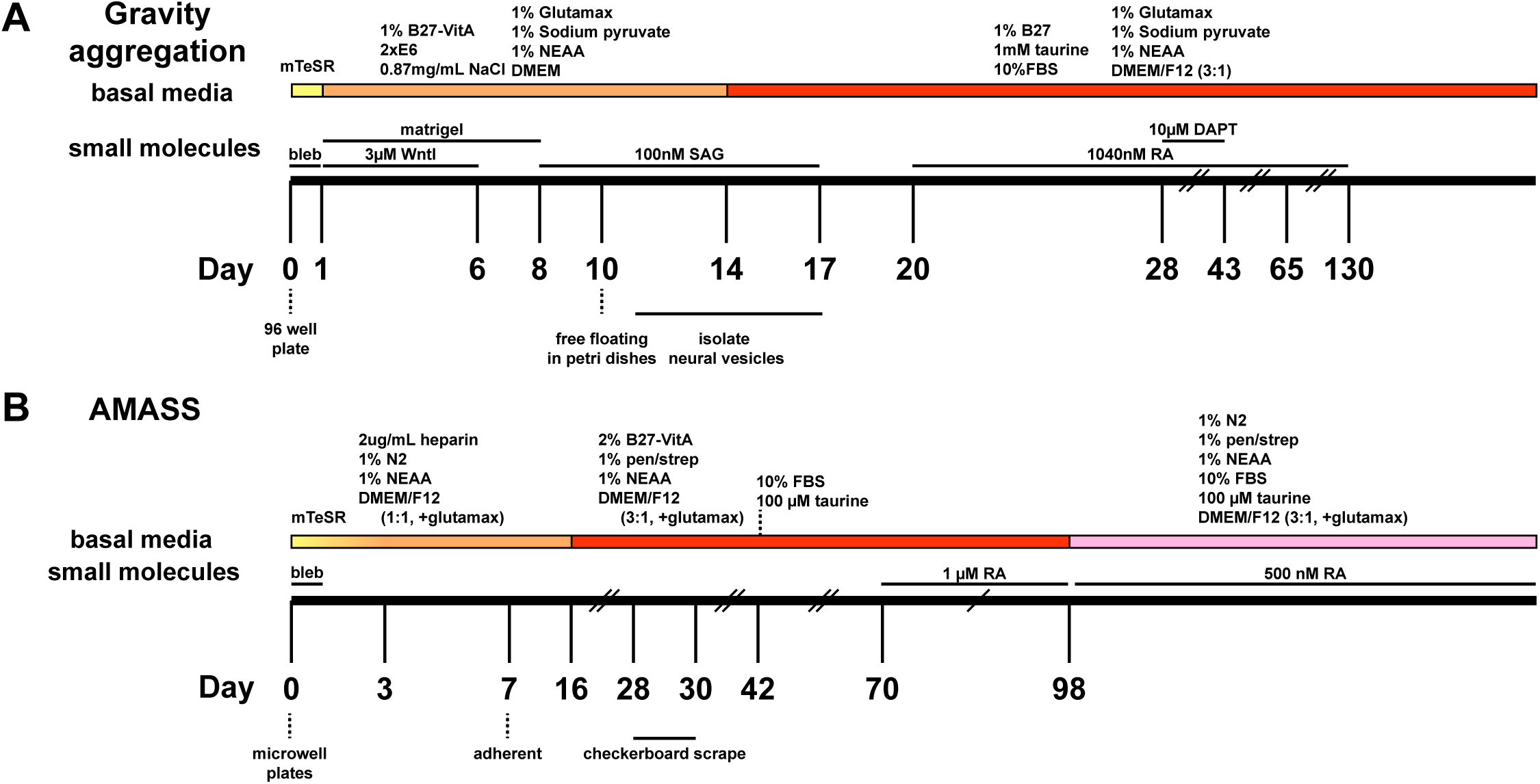
Methods of organoid differentiation. (A) Gravity aggregation adapted from (Eldred et al., 2018; Hussey et al., 2024). (B) AMASS differentiation protocol adapted from(Cowan et al., 2020).

## Materials and methods

### Cell line maintenance

H7 ESC (WA07, WiCell), H9 ESC (WA09, WiCell) modified with CRX>tdTomato (gift from David Gamm (Phillips et al., 2018)), or IMR90-GFP iPSC cells (Wahlin et al., 2021) were used for differentiation. Cells were maintained in mTeSR1 (85850, StemCell Technologies) on 1% (vol/vol) Matrigel (354263, Corning) or Geltrex (A1413202, ThermoFisher) coated dishes and grown at 37°C in a HERAcell 150i or 160i hypoxic incubator (10% CO_2_, 5% O_2_). Cells were passaged every ∼5 days as in Eldred et al 2018 (Eldred et al., 2018). Cells were passaged with Accutase (07922, StemCell Technologies) for 7-10 minutes and dissociated to single cells through additional manual trituration. Cells suspended in Accutase were added 1:2 to a solution of mTeSR1 with 5µM blebbistatin (B0560-5MG, MilliporeSigma). Cells were pelleted at 300xg for 5 minutes before being resuspended in mTeSR1 with 5µM blebbistatin. Cells were plated on a new Matrigel-GFR or Geltrex coated 6 well plate at ∼5000-40000 cells per well. Media was replaced with mTeSR1 48 hours after passaging and every day until passaged again. To minimize cell stress, no antibiotics were used.

### Cell culture media

**Stem cell media**: mTeSR1

**E6 supplement**: 970 μg/mL Insulin (11376497001, Roche), 535 μg/mL holotransferrin (T0665, Sigma), 3.20 mg/mL L-ascorbic acid (A8960, Sigma), 0.7 μg/mL sodium selenite (S5261, Sigma).

**BE6.2 media**: 2.5% E6 supplement (above), 2% B27 supplement (50X) minus Vitamin A (12587010, Gibco), 1% Glutamax (35050061, Gibco), 1% NEAA (11140050, Gibco), 1 mM Sodium Pyruvate (11360070, Gibco), and 0.87 mg/mL NaCl in DMEM (11885084, Gibco).

**LTR (Long-term retina) media**: 25% F12 (11765062, Gibco), 2% B27 supplement (50X) (17504044, Gibco), 10% heat-inactivated FBS (16140071, Gibco), 1mM Sodium Pyruvate (11360070, Gibco), 1% NEAA (11140050, Gibco), 1% Glutamax (35050061, Gibco), and 1 mM taurine (T-8691, Sigma) in DMEM (11885084, Gibco).

**NIM (neural induction medium):** 1% N2 supplement (17502048, ThermoFisher), 1% MEM NEAA, 2µg/mL heparin (H3149, SigmaAldrich) in DMEM/F12 with GlutaMAX supplement (10565018, Gibco)

**3:1 medium**: 24% F12 with GlutaMAX (31765035, Gibco), 2% B27 supplement minus Vitamin A, 1% MEM NEAA, 1% penicillin streptomycin in DMEM with high glucose, GlutaMAX, pyruvate (10569010, Gibco)

**3:1* medium**: 22% F12 with GlutaMAX, 2% B27 supplement minus Vitamin A, 1% MEM NEAA, 1% penicillin streptomycin, 10% heat inactivated FBS, 100µM taurine in DMEM with high glucose, GlutaMAX, pyruvate

**N2 medium:** 24% F12 with GlutaMAX, 1% N2 supplement, 1% MEM NEAA, 1% penicillin streptomycin, 10% heat inactivated FBS, 100µM taurine, in DMEM with high glucose, GlutaMAX, pyruvate

**Retinoic acid treatment:** Stocks of 10.4 mM retinoic acid (ATRA; R2625, Sigma) were made in DMSO. Retinoic acid was added to media for final concentrations of 1.04µM or 0.52µM.

**Thyroid hormone treatment**: 20 nM T3 (T6397, Sigma) in LTR as used previously in mouse retinal explant culture (Roberts et al., 2006) and human retinal organoids (Eldred et al., 2018). In +T3 conditions, T3 was added from day 42 until collection for analysis.

### Organoid differentiation (Fig. S7)

Organoids were differentiated from H7 WA07, H9 WA09 ESCs or IMR90 Six6>GFP iPSCs. hPSC cultures were monitored for spontaneous differentiation and only well-maintained cultures were used for differentiation. Organoids were regularly monitored for health and were culled if retinal tissue was not apparent.

### Gravity aggregation (GA)

GA differentiation proceeded as described in (Eldred et al., 2018; Hussey et al., 2024; Wahlin et al., 2017) (Fig. S7A). Cells were passaged in Accutase at 37C for 10-12 minutes and triturated to ensure single-cell dissociation. Cells were seeded in 50 µl of mTeSR1 with 5µM blebbistatin at 3,000 cells per well in 96-well ultra-low adhesion round-bottom Lipidure-coated plates (#7007, Corning). Cells were placed in hypoxia (10% CO_2_, 5% O_2_) at 37°C for 24 hours during which time cells aggregated by gravity.

After 24 hours and for the remainder of the differentiation, aggregates were grown in normoxic conditions (5% CO_2_) at 37°C. On days 1-3, 50µL of Be6.2 media with 3µM Wnt inhibitor (681669, EMD Millipore) and 1% (v/v) Matrigel were added to each well. On days 4-9, 100µL of media were removed from each well and replaced with 100µL of fresh media. On days 4-5 were Be6.2 media with 3µM Wnt inhibitor and 1% (v/v) Matrigel were added, and on days 6-7 Be6.2 media with 1% (v/v) Matrigel were added. On days 8-9 media was replaced with Be6.2 with 1% Matrigel and 100 nM smoothened agonist (SAG; 566660-1MG, SigmaAldrich).

On day 10, aggregates were rinsed 2-3x in DMEM and transferred to 10cm petri dishes in Be6.2 in 100 nM SAG. For the remainder of the differentiation, media was changed every other day. On days 13-16, aggregates were transferred to LTR media with 100 nM SAG.

Between days 11 and 16 aggregates were dissected with tungsten needles as needed to isolate optic vesicles. To remove dead cells, aggregates were washed 2-3X in DMEM as needed before feedings between days 16 and 50.

On days 16-20, aggregates were maintained in LTR. From days 20-130, 1 µM all-trans retinoic acid (RA) was added to LTR. Days 28-42 were additionally supplemented with 10 µM gamma secretase inhibitor (DAPT; 565770-10mg, CalBiochem). Organoids were maintained in unsupplemented LTR from days 131-200.

### AMASS

AMASS differentiation was described in (Cowan et al., 2020) (Fig. S7B). AggreWell 800 plates (34811, StemCell Technologies) were prepared day-of differentiation as per manufacturer’s directions. Wells were washed once with mTeSR1. ESCs were dissociated to single cell in Accutase with additional manual trituration. ESCs were resuspended to desired density in 0.5mL and this suspension was added to 1mL mTeSR1+5µM blebbistatin per well of the microwell plate and returned to hypoxia at 37°C. To generate chimeras, the desired percentage of each genotype was calculated and appropriate amounts of cells were added to reach a consistent total density of cells in one 0.5mL well.

On day 1, aggregates received a 1/3 media exchange with NIM. Day 2 received a 1/2 media replacement with NIM. Days 3-6 received a full media exchange with NIM. On day 7, aggregates were transferred in NIM to a Geltrex-coated 6 well plate.

On days 8-15 the adherent aggregates were fed daily with full media replacement with NIM. Days 16-27 were fed with 3:1 media.

Plates were scraped on day 28. Aggregates were washed and transferred to 10cm petri dishes in 3:1 media. From day 28 until the end of the differentiation, organoids were fed 3x/week in 10cm petri dishes.

Organoids were fed with 3:1 media from day 28-day 41, 3:1* from day 42-97, and N2 from day 98 until the end of differentiation. Media was supplemented with 1µM RA from days 70-97 and 0.5µM RA from day 98 until the end of differentiation. +T3 conditions were treated with T3 starting at day 42 and persisted until collection.

### Mycoplasma monitoring

Cell lines and organoid media were tested monthly for mycoplasma using MycoAlert (LT07, Lonza) and excluded if positive.

### CRISPR/Cas9 Mutations

CRISPR was performed as previously described in (Eldred et al., 2018; Hussey et al., 2024). Cloning gRNA plasmids gRNA transfection plasmids were created using pSpCas9(BB)-P2A-Puro plasmid modified from the pX459_V2.0 plasmid (62988, Addgene) by replacing T2A with P2A. gRNAs were cloned into the modified vector following the Zhang lab protocol: https://pharm.ucsf.edu/sites/pharm.ucsf.edu/files/xinchen/media-browser/CRISPR%20cloning%20protocol%20Zhang%20Lab.pdf

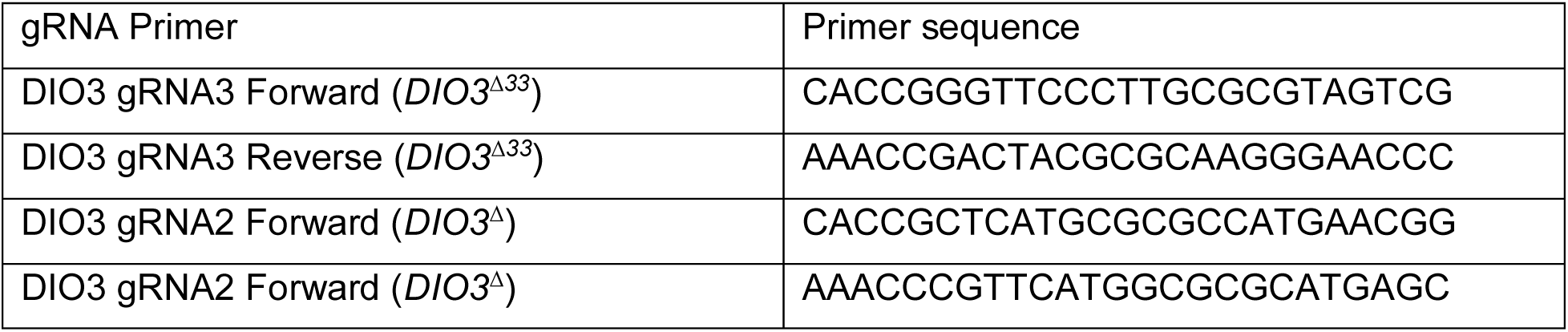

Transfection, selection, and genotyping: H7 ESCs were passaged to single cell and seeded at a density of 50,000 cells/well in mTeSR1 with 5µM blebbistatin in a prepared Geltrex or Matrigel coated 24 well plate and left in hypoxia at 37°C. 24 hours after seeding, media was replaced with mTeSR1. The transfection mix was prepared by combining: 200ng of either modified pX459 plasmid (containing gRNA) or pX458 (Cas9>GFP positive control), 50µL Optimem (31985070, Gibco) and 2µL Lipofectamine Stem Reagent (STEM00003, Invitrogen), prepared according to manufacturer’s recommendations. Each transfection well received 50µL of transfection mix.

24 hours later, media was replaced with mTeSR1. A gradient of puromycin concentrations was applied to the plate ranging from 0.1-1.5µg. After 24 hours, cells were washed once with mTeSR1 then fed with fresh mTeSR1. At this time, transfection efficiency was estimated by observing GFP+ colonies in the Cas9>GFP transfected control cells. After several days the condition in which the gRNA transfected cells survived puromycin treatment but the control Cas9>GFP transfected cells did not was passaged and expanded into a new Geltrex-coated 6 well plate. Individual colonies were manually isolated and split for either genotyping or maintenance in a new Geltrex-coated 48 well plate. DNA was extracted using QE buffer (QE0905T, Lucigen) and PCR was used to amplify the relevant region of the gene. Mutations were identified by Sanger sequencing.

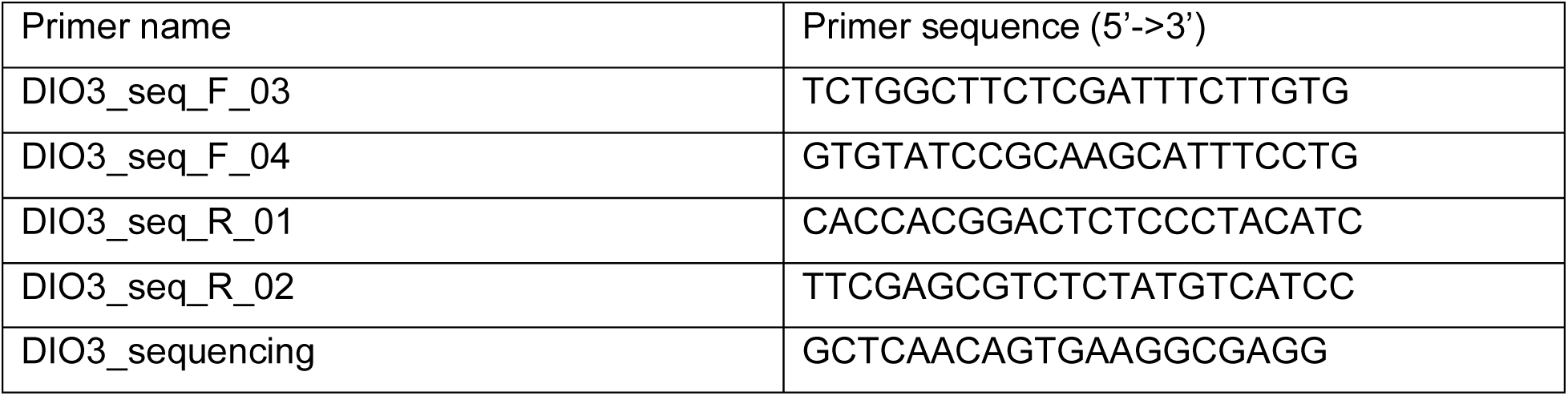

### Immunohistochemistry

Retinal organoids were fixed in freshly prepared 4% formaldehyde in PBS for 50 minutes – 1.25 hours at 4°C. Organoids were washed 3x in 5% sucrose in PBS, then transitioned through a sucrose gradient of: 6.25%, 12.5%, and 25% sucrose in PBS. All sucrose incubations were performed at 4°C and were 20 minutes – 1 hour long, with the exception of the 25% sucrose which was incubated overnight.

*Wholemount*: Organoids were transferred from 25% sucrose to block solution (2-5% donkey serum, 0.2-0.3% TritonX-100 in PBS) and incubated at 4°C for 2-120 hours. Organoids were incubated in primary antibodies diluted in block for 16-72 hours at 4°C. Organoids were washed 3x for >15 min at RT in PBS then incubated in secondary antibodies diluted in block solution for 4-48 hours at 4°C. Organoids were washed with 3x >10 min incubations in PBS at RT. If organoids were stained with Hoechst, they were incubated for 10 minutes in Hoechst 33342 (Biotium 40046 #10H0212) diluted 1:2000 in PBS at RT, followed by 3x PBS washes. Organoids were mounted in Slowfade.

*Cryosections*: Following the sucrose gradient, organoids were flash frozen in OCT and stored long term at −80°C. Organoids were cryosectioned at 10-12µm. Sections were left to dry at RT 6 hours - overnight, then stored long term at −80°C. Sections were allowed to come to RT, then rehydrated 10 min in PBS. Slides were incubated for 15-30 minutes at 60°C before proceeding with IHC. Block, primary, and secondary incubations were performed as described for wholemount organoids.

*Fetal tissue*: 14 week old fetal eyes (male, left/right eye information unknown) were enucleated within 24 hours of termination, flash frozen in liquid nitrogen, then stored at −80 °C for downstream processing. Fetal eyes were briefly thawed in RT 1x PBS, then transferred to 10% NBF and fixed overnight at 4°C. The retina was dissected from the whole globe and transferred to block solution. IHC was performed as described for organoid wholemount tissue.

### Antibodies

| Antibody | Source | Host | Dilution |
| --- | --- | --- | --- |
| ARR3 | NBP1-37003 | Goat | 1:200 |
| CRX | UJOHO013 | Mouse | 1:500 |
| CRX | WH0001406M2 | Mouse | 1:200 |
| DIO3 | NBP1-05767 | Rabbit | 1:200 |
| Ki67 | sc-23900 | mouse | 1:200 |
| NRL | AF2945 | Goat | 1:200 |
| OPN1LW/OPN1MW | AB5405 | Rabbit | 1:200 |
| OPN1LW/OPN1MW | Jeremy Nathans | Chick | 1:200 |
| OPN1SW | Sc-14363 | Goat | 1:200 |
| OPN1SW | Jeremy Nathans | Chick | 1:200 |
| RCVRN | AB5585 | Rabbit | 1:200 |
| RFP | 5F8-150 | Rat | 1:400 |
| Rhodopsin | GTX23267 | Mouse | 1:500 |
| Rhodopsin | MA5-11741 | Mouse | 1:500 |
| VSX2 | X1180P | Sheep | 1:100 |

All secondary antibodies were raised in donkey, Alexa Fluor conjugated, and used at 1:400 dilution.

### Microscopy and Image Processing

Brightfield images were acquired with either a Zeiss LSM 980 inverted microscope with an Axiocam 506 color camera system and a C Apo 40×/1.2 W DICIII lens or an EVOS XL Core Cell imaging system on DIC with a 4x, 10x, or 20x air objective lens (Thermo Fisher).

Fluorescent images were acquired using either a Zeiss LSM 800 or 980 laser scanning confocal, or a Nikon CSU-W1 SoRa spinning disk system. Confocal images were acquired with similar settings for laser power, photomultiplier gain and offset, and pinhole diameter, and camera images were acquired at similar epifluorescence intensity and exposure. Cones were imaged at 30-100 optical sections, 1.26µm step size for confocal or 0.50 µm for camera, and visualized with a maximum intensity projection or a single Z section. For quantitative measurements of signal intensity, tissues were imaged at the same imaging parameters in the same imaging session to limit variability.

### Measurements and Quantification

Manual quantification of photoreceptors was performed with Adobe Photoshop and density was calculated using ImageJ. Semi-automated quantification of photoreceptors was done using the Object Colocalization FL v2.1.4 module of HALO Image Analysis Platform 3.5.3577 (Indica Labs) used to count the number of S-, L/M-opsin, and Rhodopsin-expressing cells. Outer segment length was quantified using Fiji in a single Z that captured the focal plane of the OS. DIO3 fluorescence intensity was analyzed in Fiji using a single z that captured the center focal plane of the cells being analyzed. 50 cells were selected per clone type per organoid that were in plane and CRX+. An appropriate radius was selected to encircle each cell and the integrated density and standard deviation of the population was recorded.

GraphPad Prism was used for analysis and statistics. Statistical tests and N are listed in figure legends. All error bars represent SEM. Organoids with fewer than 150 cones after day 120 were removed from analysis and assumed to not have properly differentiated.

### DIO3 enzymatic activity assay

Type 3 deiodinase activity was determined as previously described (Hernandez et al., 2006). In brief, organoids were homogenized in 100 µL of Tris 10 mM/Sucrose 0.25M pH=7.4 in the presence of 2 mM dithiothreitol (DTT). 10 µL of the homogenate was assayed for protein content. For the enzymatic assay, 5 µL of the organoid homogenate were incubated with 100 fmols of radiolabeled T3 for half an hour at 37.5 °C. The incubation mix contained 20 mM DTT and 1 mM propylthiouracyl in a total volume of 50 µL. The reaction was stopped by adding 10 µl of 5 µM T3 and 50 µL of ethanol. An aliquot of the reaction was submitted to paper chromatography as described (Hernandez et al., 2006), and radiolabeled T3 and T2 (3,3’-diiodothyronine) were quantified in a gamma counter to calculate the proportion of T3 deiodinated and enzymatic activity.

### Nuclei Lysis

Nuclei were isolated from retinal organoids using a lysis protocol previously described (Santiago et al., 2023). In brief, 5 retinal organoids per genotype were pooled and flash frozen and stored at −80°C until the nuclei were extracted using chilled lysis buffer (10 mM Tris-HCl, 10 mM NaCl, 3 mM MgCl_2_, 0.01% Tween-20, 0.01% Nonidet P-40, 0.001% Digitonin, 1mM DTT, 1% BSA). The homogenate was incubated on ice for 10 minutes, washed twice with a resuspension buffer (10 mM Tris-HCl, 10 mM NaCl, 3 mM MgCl_2_, 0.1% Tween-20, 1mM DTT, 1% BSA) and filtered through a series of 70 μm and 40 μm Flowmi filters. The nuclei were resuspended in an appropriate volume of diluted nuclei buffer (10X Genomics) to achieve a concentration of approximately 2000–5000 nuclei/μL. Nuclei concentration was determined using trypan blue and DAPI staining.

### Single-nucleus multiomics Library Construction and Sequencing

Single-nucleus RNA-sequencing and ATAC-seq were performed on dissociated retinal nuclei using the Chromium Single Cell Multiome ATAC + Gene Expression Reagent Kits (10× Genomics, Pleasanton, CA). Briefly, nuclei (∼16000 nuclei per sample) were processed for tagmentation and loaded onto the 10X Chromium controller and the downstream RNA or ATAC libraries were generated and indexed by following the manufacturer’s instructions. Libraries were pooled and sequenced on Illumina NovaSeq 6000 targeting 50000 reads per cell for each RNA and ATAC library.

### Single-cell muti-omics analysis

The sequencing data from the single cell RNA and ATAC libraries were demultiplexed and processed using Cell Ranger ARC v2.0.2 (10X Genomics). Reads were mapped to the GRCh38 reference genome and count matrices were generated. Filtered feature-barcode matrices were loaded for analysis using the Seurat and Signac packages in R (Hao et al., 2021; Stuart et al., 2021). Initial quality control was performed on the RNA assay to filter low-quality cells and doublets were identified using the scDblFinder R package (Germain et al., 2021). Nuclei with more than 3% of reads mapped to mitochondrial genes, fewer than 900 unique molecular identifiers (UMIs), or more than 50,000 UMIs were excluded. Datasets were merged and underwent normalization, variable feature selection and scaling using Seurat’s *NormalizeData*, *FindVariableFeatures*, and *ScaleData* functions. Principal components were then calculated and batch corrected using the Harmony R package (Korsunsky et al., 2019). Uniform Manifold Approximation and Projection (UMAP) dimension reduction was performed on the corrected principal components, and clusters were computed using Seurat’s *FindNeighbors* and *FindClusters* functions. Cell types were then identified using a list of known marker genes that were used previously (Liu et al., 2023). ATAC peaks were called using MACS2 to generate peak sets for each cell type and then merged to create a consensus peak set, which was quantified for each cell (Zhang et al., 2008). Low-quality peaks and regions overlapping blacklisted genomic regions were filtered out. Further filtering on the data was performed with nuclei fewer than 1000 or greater than 50,000 ATAC fragments were removed. Additionally, nuclei were retained if they had a nucleosome signal less than 2 and a TSS enrichment score greater than 1. Latent semantic indexing (LSI) was applied to reduce the dimensionality of the peak matrix, followed by batch correction using Harmony. Gene activity scores and peak-to-gene links were calculated using Signac’s *GeneActivity* and *LinkPeaks* functions. Motif analysis using the JASPAR2020 core vertebrates collection was performed and ChromVAR was employed to calculate motif accessibility deviations per cell (Fornes et al., 2020; Schep et al., 2017). Cell trajectories were calculated using the Slingshot R package and the pseudotime estimates were used to visualize the progression of cells along inferred lineages (Street et al., 2018). The UCell R package was used to calculate enrichment scores for S cones, L/M cones and Rods using a curated set of genes published previously (Andreatta and Carmona, 2021; Hussey et al., 2024).

### Computational modeling

We simulated our mathematical model of photoreceptor specification using a hybrid deterministic-stochastic approach implemented in MATLAB. The concentration of T3—due to large numbers of individual molecules even at pM concentrations—was simulated deterministically via numerical integration. All other species in the model were simulated stochastically using the Gillespie algorithm. All model equations, stochastic events, rate functions and parameters are detailed in the supplementary information. All curves in **Fig. 6B** are single representative simulations of organoids seeded with 10^4^ RPCs. Simulations were run until each cell in the system adopted a terminal fate. The histograms in **Fig. 6C** compile the results of 5000 replicate simulations, each of which was seeded with a randomly determined number of initial RPC cells. Initial RPC numbers were drawn from a normal distribution with mean 10^4^ and standard deviation 2000. To ensure that the signaling and intrinsic model implementations were comparable, we adjusted the maximal rates of *S* and *L/M* photoreceptor specification in the intrinsic model (𝜆_𝑆_ and 𝜆_𝐿_) such that the final average number of each photoreceptor matched the signaling model. In **Fig. 6D**, each point summarizes the output of 5000 replicate simulations, each of which was initiated with a random number of RPCs as in **Fig. 6C**. Each set of simulations was run with a randomly seeded parameter set. Parameters were logarithmically sampled over a 100-fold range.

## Data and Code Availability

All sequencing data generated in this study can be accessed at GEO accession number GSE292057 (*wildtype* vs *DIO3^Δ33^* mutant organoid single nucleus multiomics). Code used to analyze the single cell datasets in this study can be found at https://github.com/csanti88/dio3_christina. *wildtype* vs *wildtype* +T3 treated organoid single cell RNA sequencing data was generated in (Hussey et al., 2024)

## A stochastic model of T3-regulated photoreceptor specification in retinal organoids

Here we detail the computational model used to simulate thyroid hormone-mediated control of retinal organoid development presented in **Fig. 6** of the main text. This document will walk through model construction (Section 1: ‘*Model Construction’*), build intuition for the dynamics of cell fate specification in wild-type and *DIO3* mutant organoids (Section 2: ‘*Simulation of Cell Fate Specification Dynamics’*), and present a speculative discussion of how thyroid hormone regulation could be used to achieve robust organoid development (Section 3: ‘*Robustness in Retinal Organoid Development*).

### Model Construction

#### 1.1. Structure and key assumptions

In our model, feedback control over photoreceptor specification is exerted through the regulation of thyroid hormone abundance. Specifically, retinal progenitor cells (*P*) express DIO3, an enzyme that degrades active thyroid hormone (*T*). As organoid development proceeds, DIO3-expressing progenitors are depleted, resulting in an increasing concentration of *T* and acceleration of photoreceptor specification. As we will demonstrate, these features are sufficient to reproduce the observed photoreceptor specification dynamics of wild-type and *DIO3* mutant organoids and suggest that thyroid hormone-mediated feedback could be used to suppress noise in photoreceptor specification.

The model makes the following mechanistic assumptions:

1. Thyroid hormone (*T*) induces the differentiation of L/M-opsin-expressing (*L*) and S-opsin-expressing (*S*) photoreceptor cells from retinal progenitor cells (*P*).
2. *P* cells can also differentiate into non-cone fates (collectively referred to as *A* cells) in a thyroid-independent manner.
3. Terminal commitment to *S* and *L* fates is not instantaneous. Immature *S* and *L* cells (*S_im_* and *L_im_* cells, respectively) are generated from *P* cells, and irreversibly commit to mature fates with a constant rate (𝜆_𝑚_).
4. Immature photoreceptor cells can enter into a ‘double expressing’ state (*SL*) in which both *L/M* and *S* opsins are expressed. These cells probabilistically mature into terminal *S* or *L* cells with constant, *T*-independent per-cell rates.
5. *T* is imported into the organoid at a constant rate (𝜆_𝑇_) and is degraded by the action of DIO3. Our measurements of intra-organoid *T* concentration (< 60 pM) are two orders of magnitude below the empirical K_m_ of DIO3 (Kuiper et al., 2003). We therefore assume that the rate of DIO3-mediated T3 degradation is linear with respect to T3.
6. DIO3 is expressed in *P* cells, but not in differentiated photoreceptor cells.
7. Rates of *S_im_* and *L_im_* differentiation are saturable in *T*. For simplicity, we relate *T* to the rate of differentiation with a Hill function (Hill constant = 1).

The flow of cells through available states and their accompanying rates of each event can be graphically summarized as follows:

**Figure 1.1.**
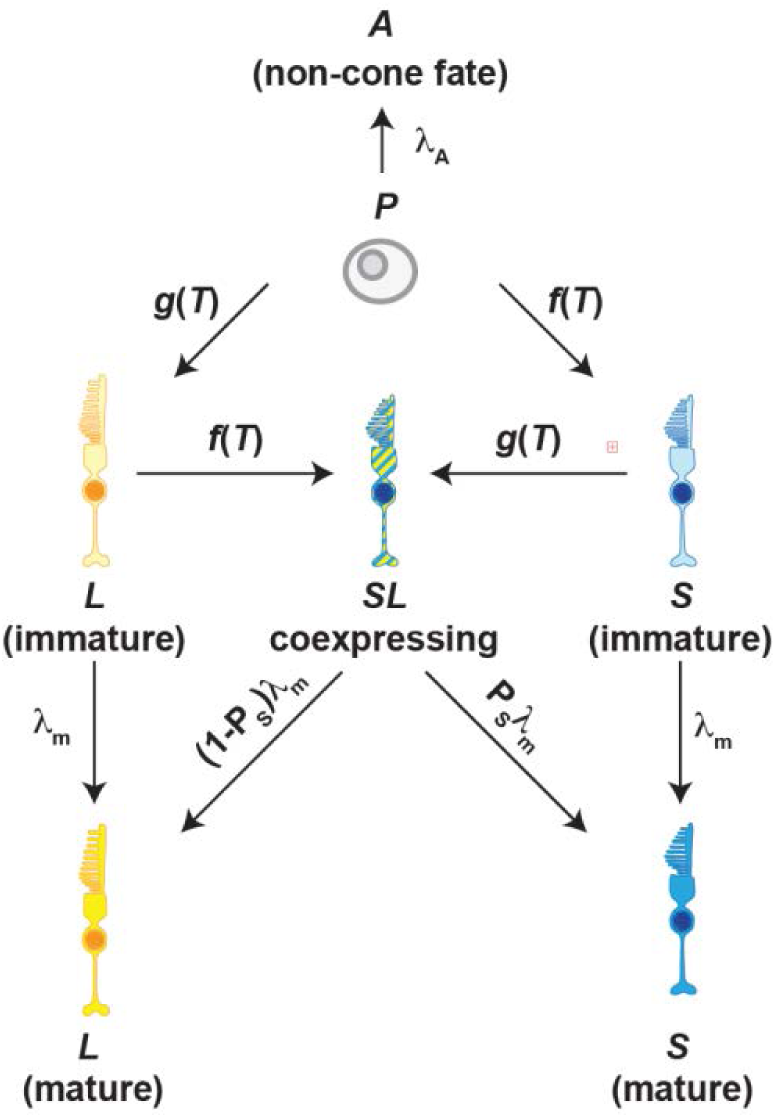
Model schematic. In each simulation, all cells begin as undifferentiated *P* cells and can proceed along differentiation trajectories as summarized by the arrows. The expression above each arrow denotes the form of the per-cell rate of transition along that pathway. Simulations conclude when all cells have adopted terminal fates (i.e. *L, S,* or *A*).

We simulate cell differentiation dynamics as a stochastic birth-death process. All simulations presented in the main text were carried out using a modified Gillespie algorithm in which *T* dynamics were simulated deterministically via numerical integration and cell fate decisions occur stochastically. Below, we formalize each permissible event, its state-dependent rate and the resulting step sizes for each system component.

Deterministic components:

1. **T3 Dynamics**: 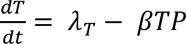

Stochastic components:

1. Immature *S* Birth: 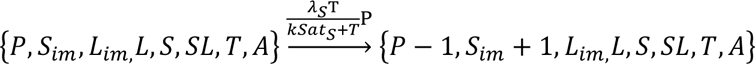
2. Immature *L* Birth: 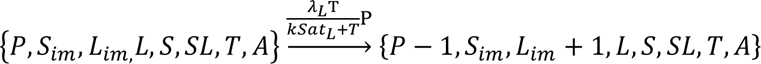
3. *SL* Birth (from *S_im_*): 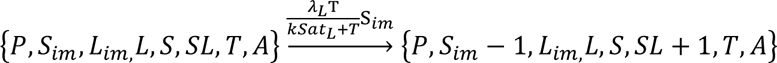
4. *SL* Birth (from *L_im_*): 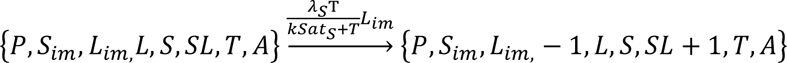
5. *S* maturation: 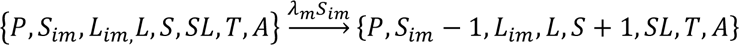
6. *L* maturation: 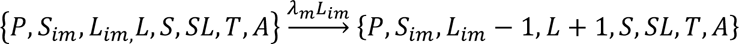
7. *SL* matures to *S*: 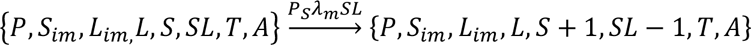
8. *SL* matures to *L*: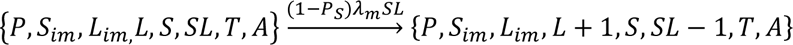
9. *A* birth: 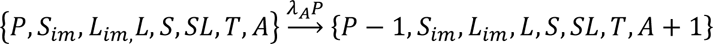

#### 1.2. Event Rates

In the table below, we summarize the rationale behind the form of each event rate.

| Process | Form | Intuition |
| --- | --- | --- |
| $T$ import | $\lambda_T$ | We assume that $T$ is transported into the organoid at a constant rate. Note that $T$ import is simulated deterministically. |
| $T$ degradation | $\beta TP$ | $T$ is degraded with first-order kinetics by DIO3-expressing $P$ cells. The effective rate constant of degradation is the per-cell rate of $T$ degradation ($\beta$) multiplied by the number of degrading cells ($P$). Note that $T$ degradation is simulated deterministically. |
| $S_{im}$ Birth | $\frac{\lambda_S T}{kSat_S + T} P$ | The per-cell rate of $P \rightarrow S_{im}$ differentiation ($f(T)$ in diagram above) is a saturating function of $T$ for which the half-maximal value ($\lambda_S/2$) occurs at $T = kSat_S$. |
| $L_{im}$ Birth | $\frac{\lambda_L T}{kSat_L + T} P$ | The per-cell rate of $P \rightarrow L_{im}$ differentiation ($g(T)$ in diagram above) is a saturating function of $T$ for which the half-maximal value ($\lambda_L/2$) occurs at $T = kSat_L$. |
| $SL$ Birth (from $S_{im}$) | $\frac{\lambda_L T}{kSat_L + T} S_{im}$ | The per-cell rate of $S_{im} \rightarrow SL$ differentiation ($g(T)$ in diagram above) is a saturating function of $T$ for which the half-maximal value ($\lambda_L/2$) occurs at $T = kSat_L$.<br>We note that this is identical to the rate of $P \rightarrow L_{im}$ differentiation. Intuitively, this reflects an assumption |
| | | that expression of $L$-related genes is independent of adoption of the $S_{im}$ fate. |
| $SL$ Birth (from $L_{im}$) | $\frac{\lambda_B T}{kSat_B + T} L_{im}$ | The per-cell rate of $L_{im} \rightarrow SL$ differentiation ($f(T)$ in diagram above) is a saturating function of $T$ for which the half-maximal value ($\lambda_L/2$) occurs at $T = kSat_L$.<br>We note that this is identical to the rate of $P \rightarrow S_{im}$ differentiation. Intuitively, this reflects an assumption that expression of $S$-related genes is independent of adoption of the $L_{im}$ fate. |
| $S_{im}$ maturation | $\lambda_m S_{im}$ | Maturation occurs with a constant rate ($\lambda_m$) per cell. |
| $L_{im}$ maturation | $\lambda_m L_{im}$ | Maturation occurs with a constant rate ($\lambda_m$) per cell. |
| $SL$ matures to $S$ | $P_S \lambda_m SL$ | <u>$SL$</u> Maturation occurs with the same constant rate ($\lambda_m$) per cell as with $S_{im}$ and $L_{im}$, but probabilistically resolves into a mature $S$ cell with probability $P_S$. |
| $SL$ matures to $L$ | $(1 - P_S) \lambda_m SL$ | <u>$SL$</u> Maturation occurs with the same constant rate ($\lambda_m$) per cell as with $S_{im}$ and $L_{im}$, but probabilistically resolves into a mature $L$ cell with probability $1 - P_S$. |
| $A$ birth | $\lambda_A P$ | Progenitor cells adopt a non-cone alternative fate with a constant, $T$-independent rate per cell ($\lambda_A$). |

#### 1.3. Model Parameter Values

| Parameter | Description | Value | Intuition for value |
| --- | --- | --- | --- |
| $\lambda_T$ | Rate of T3 import | 24 pM/day | With $\beta$, sets initial pseudo-steady state $T$ concentration at 1 pM |
| $\beta$ | Rate constant for T3 degradation | $2.4 \times 10^{-3} \text{ day}^{-1}$ | With $\lambda_T$, sets initial pseudo-steady state $T$ concentration at 1 pM and initial $T$ average lifetime at 1 hour. |
| $\lambda_S$ | Maximal rate of $S_{im}$ | $7.2 \times 10^{-2} \text{ day}^{-1}$ | Sets average $P \rightarrow S_{im}$ waiting time at 14 |
| | differentiation from $P$ | | days at saturating $T$ concentrations. |
| $kSat_S$ | T3 concentration at which rate of $S_{im}$ differentiation is half-maximal | 5 pM | Rate of $P \rightarrow S_{im}$ differentiation is half maximal when $P$ decreases to 2000 cells. |
| $\lambda_L$ | Maximal rate of $L_{im}$ differentiation from $P$ | $2.5 \times 10^{-1} \text{ day}^{-1}$ | Sets average $P \rightarrow L_{im}$ waiting time at 4 days at saturating $T$ concentrations. |
| $kSat_L$ | T3 concentration at which rate of $L_{im}$ differentiation is half-maximal | 40 pM | Rate of $P \rightarrow L_{im}$ differentiation is half maximal when $P$ decreases to 250 cells. |
| $\lambda_A$ | Rate of $A$ differentiation from $P$ | $2.5 \times 10^{-2} \text{ day}^{-1}$ | Sets average $P \rightarrow A$ waiting time at 40 days. |
| $\lambda_{mat}$ | Maturation rate for $L_{im}$, $S_{im}$ and $SL$ cells. | $6.1 \times 10^{-2} \text{ day}^{-1}$ | Average residence time in the immature state is 16 days. |
| $P_S$ | Probability of a $SL$ co-expressing cell resolving into a mature $S$ cell. | 0.1 | 90% of $SL$ cells resolve into $L$ mature state. |

### Simulation of Cell Fate Specification Dynamics

The model aims to capture several observations presented in the main text. Specifically, it was designed to accommodate the following:

1. *S* and *L/M* cone specification is induced by thyroid hormone signaling.
2. In *wildtype* organoids, *S* cone specification precedes *L/M* cone specification, but the *L/M* population ‘overtakes’ the *S* population late in development.
3. *DIO3* mutant organoids exhibit accelerated photoreceptor specification, increased proportion of *L/M* cones, and transient appearance of *SL* co-expressing cells.
4. Addition of exogenous *T* moderately accelerates specification dynamics in *wildtype* organoids, but leaves *DIO3^Δ33^* mutant dynamics unchanged.

Below, we reproduce the simulations from **Fig. 6** of the main text to discuss model performance with respect to these benchmarks.

#### 2.1. *wildtype* dynamics

**Figure 2.1.**
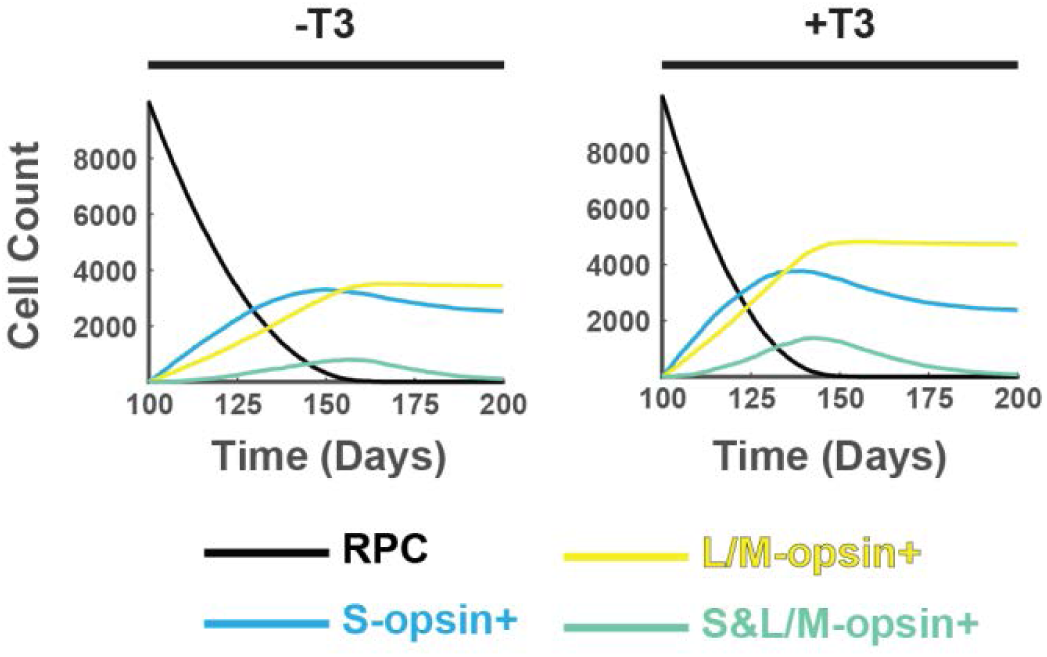
Simulated photoreceptor specification dynamics in *wildtype* organoids. The figure reproduces the wild-type panels from **Fig. 6B** of the main text. The left simulation represents unperturbed *wildtype* development, and the right represents a *wildtype* organoid grown with exogenous T3 added to the culture medium. Black, blue, yellow and green curves depict *P, S, L* and *SL* cell type copy numbers, respectively. Each simulation begins with 10^4^ undifferentiated *P* cells.

The ‘*-*T3’ simulation above (**Fig. 2.1**, left panel) captures the key features of photoreceptor specification dynamics in unperturbed *wildtype* organoids. Specifically, *S* cones appear first, with abundance peaking at approximately day 140. *L/M* cones appear later and ultimately overtake *S* in abundance. The model also predicts the observation of a small number (∼15% of the total pool) of cones that co-express *S* and *L/M* opsins. Finally, the model captures the modest acceleration of specification dynamics observed when exogenous T3 is added to culture medium (**Fig. 2.1**, right panel).

The characteristic dynamics of photoreceptor specification in *wildtype* organoids can be understood intuitively in terms of *T-*dependent model rates. *S* cells appear before *L* cells because they are induced by lower concentrations of *T* (i.e. 𝑘𝑆𝑎𝑡_𝑆_ < 𝑘𝑆𝑎𝑡_𝐿_)*. L* cells ultimately ‘catch up’ to *S* due to two factors. First, the *maximal* rate of *L* differentiation at the high *T* concentrations that occur late in development is greater than the corresponding rate for *S* cells (i.e. 𝜆_𝐿_ > 𝜆_𝑆_). Second, the *SL* co-expressing cells preferentially resolve into mature *L* cells (i.e. 𝑃_𝑆_ = 0.1). This has the net effect of ‘redirecting’ immature *S* cells into *L* fates via the *SL* co-expressing state. This second effect also explains why *S* cell numbers slightly decline after their peak.

Our model also offers a simple explanation for the existence of *SL* co-expressing cells. We posit that immature *S* and *L* cells retain the ability to ‘fire’ a second *T-*dependent developmental program—expression of *L/M* or *S* opsins, respectively— until they mature into terminally differentiated photoreceptors. Co-expressing cells thus arise late in development when *T* concentrations are high enough that the rates of *S* or *L* program initiation compete with the rate of cone maturation. Consistent with this intuition, *SL* cell abundance is elevated by both the addition of exogenous *T3* (**Fig. 2.1**, right panel) and removal of DIO3 (see below).

#### 2.2. *DIO3^Δ33^* dynamics

**Figure 2.2.**
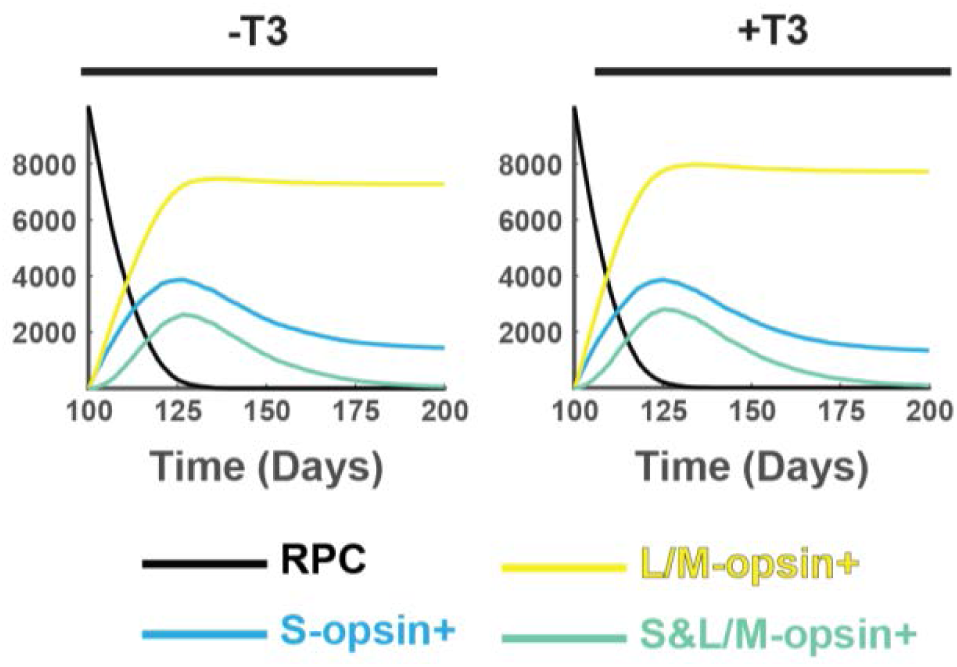
Simulated photoreceptor specification dynamics in *DIO3^Δ33^* mutant organoids. The figure reproduces the *DIO3^Δ33^* mutant panels from Fig. 6B of the main text. The left simulation represents an unperturbed *DIO3^Δ33^*mutant organoid, and the right represents *a DIO3^Δ33^* mutant organoid grown with exogenous T3 added to the culture medium. Black, blue, yellow and green curves depict *P, S, L,* and *SL* cell type copy numbers, respectively. Each simulation begins with 10^4^ undifferentiated *P* cells.

Our model reproduces the key features of photoreceptor specification dynamics in both perturbed and unperturbed *DIO3^Δ33^* mutant organoids. Specifically, it correctly captures the accelerated timeline of photoreceptor specification, increased proportion of *L* cells, as well as increased proportion of *SL* co-expressing cells. Additionally, it correctly predicts that addition of exogenous T3 has no effect on the dynamics or final composition of the photoreceptor pool.

These features can again be understood in terms of the *T-*dependence of model rate functions. The acceleration of organoid development results from increased *T* abundance; the *DIO3^Δ33^* mutant exhibits a 94% reduction in the rate constant of *T* degradation, leading to a ∼20-fold increase in *T* concentration relative to *wildtype* organoids. This results in elevated rates of *S* and *L* cell differentiation, as well as a higher abundance of *SL* co-expressing cells. This increase also explains why addition of exogenous *T* does not alter development in mutant organoids. The 20-fold increase in baseline *T* levels pushes the rates of *S* and *L* differentiation into the saturation regime, so that additional *T* has little to no effect on development.

### Robustness in Retinal Organoid Development

The material presented above demonstrates that the simple kinetic principles of our model are sufficient to reproduce the photoreceptor specification dynamics of *wildtype* and *DIO3^Δ33^* mutant organoids. In this section, we turn to the question of why T3-mediated feedback control might be useful for development. Below, we develop the hypothesis that this control confers robustness to retinal organoid development.

#### 3.1. Constructing a cell-intrinsic model

For the arguments below, it will be useful to contrast the behavior of our ‘signaling’ model with that of a ‘cell-intrinsic’ model that lacks feedback. The states and permissible transitions of the model are assumed to be identical to those laid out in **Fig. 1.1**. However, the rates with *T-* dependence become *T*-independent constants. We formalize the cell-intrinsic stochastic birth-death process as follows:

1. Immature *S* Birth: 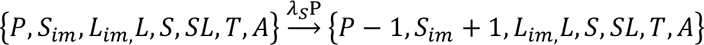
2. Immature *L* Birth: 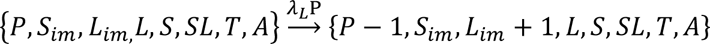
3. *SL* Birth (from *S_im_*): 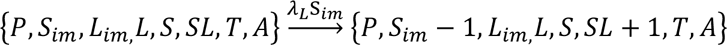
4. *SL* Birth (from *L_im_*): 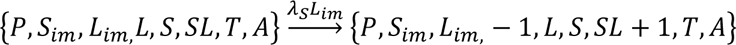
5. ***S* maturation**: 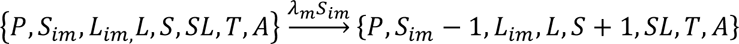
6. ***L* maturation**: 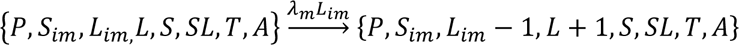
7. *SL* matures to *S*: 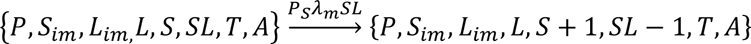
8. *SL* matures to *L*: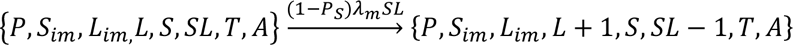
9. ***A* birth:** 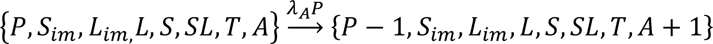

#### 3.2. Comparing robustness of signaling and cell-intrinsic models

In **Fig. 6C** and **D** of the main text, we argue that T3-mediated feedback confers robustness to noise in the initial number of *P* cells in the early organoid. This was shown for a specific set of model parameters in **Fig. 6C**; when the initial number of *P* cells in each replicate simulation is a random variable, the final number of photoreceptor cells is more tightly distributed in the signaling model than in the cell-intrinsic model. In this sense, T3 feedback enables this noise to be ‘filtered out’, leading to a more precise number of final photoreceptors across replicate realizations. Finally, we demonstrate that this property is not a product of the specific parameter sets chosen for **Fig. 6 A-C**, but rather a generic property of the feedback system across many distinct parameter sets (**Fig. 6D**, reproduced below as **Fig. 3.1**). Not only does it appear that the signaling model *can* outperform the cell-intrinsic model, but that the intrinsic model matches the *worst achievable* performance by the models with feedback. In the following sections we develop intuition for this observation.

**Figure 3.1.**
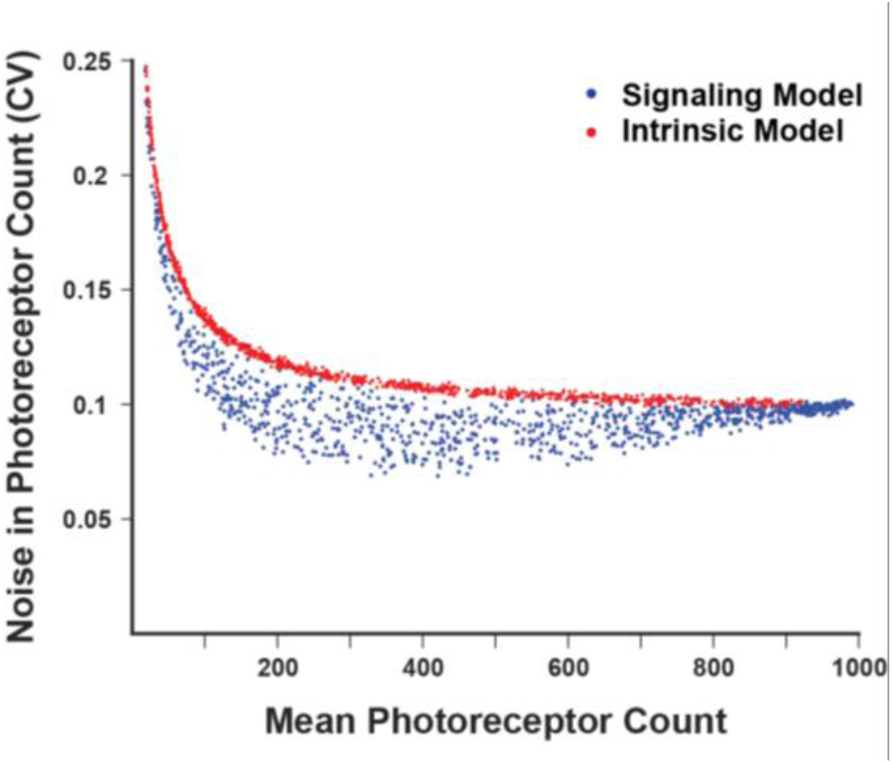
Enhanced developmental robustness in signaling models. Exploration of model parameter space. Simulations (n = 1000) were performed for signaling (blue) and cell-intrinsic (red) models with randomly sampled parameter values. Each dot represents the performance of 50 replicate simulations with a unique parameter set.

#### 3.3. Limiting behaviors of cell-intrinsic and signaling models

The empirical exploration of model capabilities presented in **Fig. 3.1** suggests the existence of two smooth, limiting curves in mean-CV space: a ‘noisy’ upper limit and a ‘precise’ lower limit that intersect at low and high mean photoreceptor counts. As we explore below, identifying the origins of these limits will help to explain how the signaling model filters out noise in initial *P* abundance.

To develop intuition for the upper limit curve shape, we recognize that the intrinsic model— from the point of view of the final photoreceptor count— can be viewed as a simple binomial cut of a randomly-varying initial number of progenitors. That is, each *P* cell present at the beginning of the simulation has only two possible end fates: cone (i.e. *S* or *L* cells) or non-cone (*A* cells, which also encapsulate the possibility of a cell death fate). Since the per-cell rates of each differentiation event constant, the probability that a given *P* cell adopts these fates are 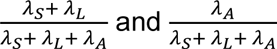, respectively. Each *P* cell makes an independent decision according to these probabilities, resulting in binomial statistics for a given initial number of progenitors. The amount of noise in photoreceptor abundance will be larger at low mean photoreceptor count— reflecting both the variation in initial *P* abundance and low-number noise due to small photoreceptor differentiation rates— and asymptotically approach the noise in *P* seeding at higher average counts when the randomness of individual fate decisions is negligible. This simplified binomial model is superimposed onto the plot below (**Fig. 3.2**, black curve), and appears to capture the upper noise limit’s shape.

The lower limit represents a diametrically-opposed kinetic regime. In the intrinsic model, the cone and non-cone differentiation rates are all first-order with respect to *P*. The lower limit can be captured, by contrast, with a simplified model in which cones and non-cones differentiate with zero- and first-order kinetics, respectively. In this limiting model, the rate of non-cone differentiation is still P-dependent (𝜆_𝐴_𝑃), while the rates of *S* and *L* differentiation are *P*-independent (i.e. 𝜆_𝑆_ and 𝜆_𝐿_, respectively). Simulating a simplified model with these zero/first-order assumptions results in a curve that appears to represent a lower bound for the behavior of the signaling model (**Fig. 3.2**, magenta curve).

**Figure 3.2.**
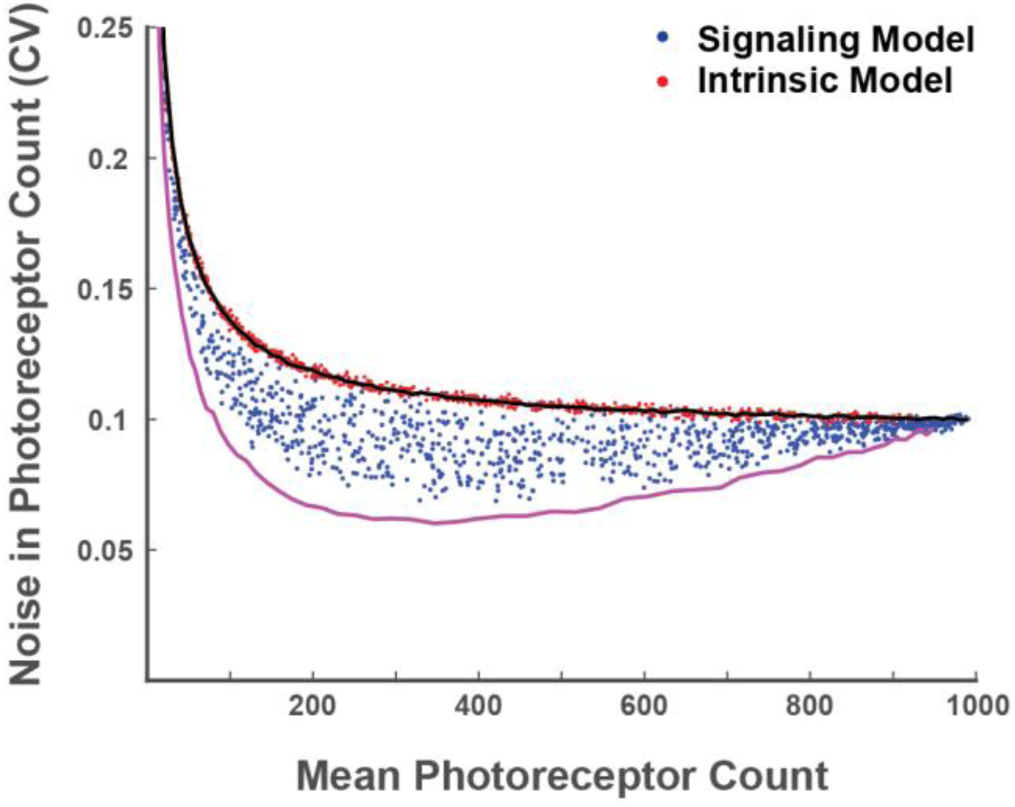
Limiting behavior of signaling and cell-intrinsic models. Blue and red dots depict simulations with randomly-sampled parameter sets as in **Fig. 3.1**. Black curve represents the binomial approximation to the cell intrinsic model. The magenta curve represents the first/zero-order approximation to the signaling model. These approximations appear to capture the limiting behaviors of the cell-intrinsic and signaling models.

#### 3.4. Intuition behind noise filtering in the signaling model

The analysis presented in the previous section demonstrates that the best-performing parameter sets in the signaling model occupy a regime where non-cone and cone differentiation rates are first- and zero-order with respect to *P*. This raises two key questions. First, why does this regime perform well? Second, how can the signaling model achieve zero-order differentiation rates for *L* and *S* cells? We address both questions in order below.

The mechanism of noise suppression in the idealized signaling model can be understood in terms of variation in the time required to consume all *P* cells. In a simple model with only zero-order differentiation of *P* cells, the variation in initial numbers of *P* translates directly into variation in the time it takes to complete the process: twice as many *P* cells take twice as long to consume when the rate of differentiation is constant (i.e. zero-order). By contrast, a model with only first-order differentiation kinetics achieves some buffering. Because the size of the *P* pool declines exponentially in this model, having twice as many initial cells results in only a log(2) increase in the completion time. Importantly, combining both first- and zero-order removal further improves this timing precision over the first-order only model. Indeed, this combination of first- and zero-order removal has been proposed to underlie the tight timing of cell fate commitments in bacterial cells (Lord et al., 2019; Norman et al., 2013).

The precise timing behavior of the signaling model is the key to the noise suppression illustrated in **Fig. 3.1**. The combination of first-order consumption of *P* to make *A* cells and zero-order consumption of *P* to make *S* and *L* cells means that the time required to complete differentiation will be similar even when the initial number of *P* cells widely varies. During this tightly-distributed time, the zero-order kinetics of cone differentiation steadily accumulate new cones in a constant, clock-like manner. These two effects together mean that the final count of cones is tightly controlled: the number of ‘ticks’ achieved by a clock over a narrowly-distributed period of time will be narrowly-distributed.

Importantly, this noise-buffering behavior is not accessible to the cell-intrinsic model because all differentiation events have first-order kinetics. This can be illustrated with a series of simulations in which the number of initial *P* cells is systematically varied. In the cell-intrinsic model, changes in the number of *P* cells translates directly into changes in the final number of photoreceptor cells (i.e. the relationship is linear, **Fig. 3.3**, red dots). By contrast, this relationship is sub-linear for the signaling molecule; an X% change in initial *P* abundance leads to a less-than X% change in final photoreceptor count (**Fig. 3.3**, blue dots). This differential sensitivity to changes in *P* underlies the noise buffering properties that we propose.

**Figure 3.3.**
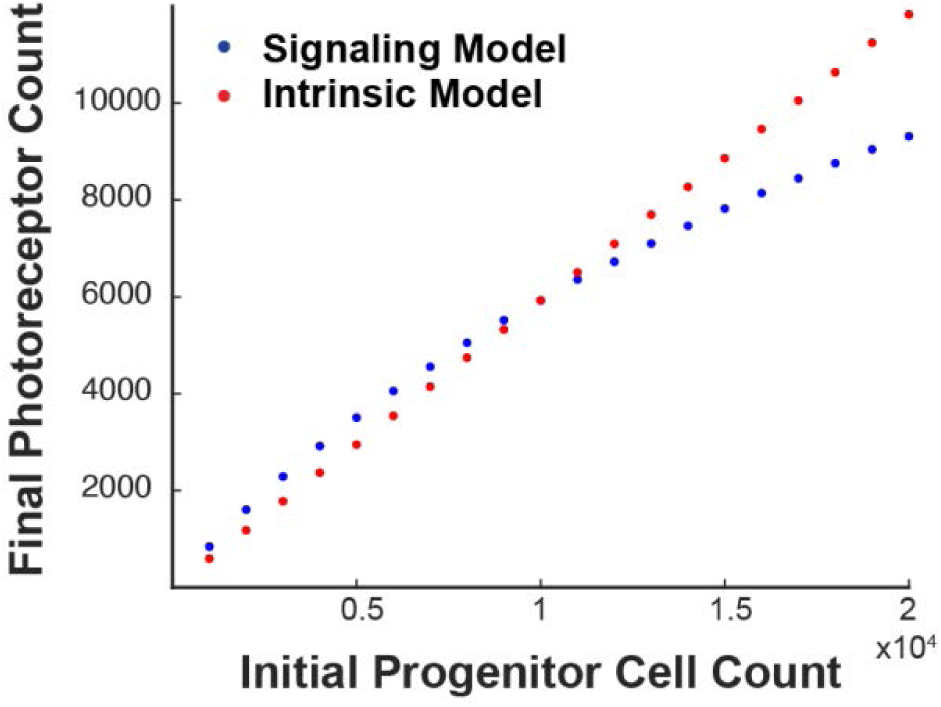
Model sensitivity to changes in initial progenitor cell abundance. This plot compares the mean number of final photoreceptors (*L* and *S* cells) to the initial number of progenitors (*P*) cells for the signaling (blue) and cell-intrinsic (red) models.

The best performing signaling molecule parameter sets mimic a situation in which *L* and *S* differentiation rates are zero-order with respect to *P*. How can this situation arise when the mathematical forms of the corresponding rate functions depend on *P*? The answer lies in the relationship between *P* and *T* abundance. *T* levels increase as development proceeds due to the depletion of *P* cells; only *P* cells degrade *T*, so thyroid hormone levels increase as the degrading cells differentiate. This relationship creates an effective cancellation: the rate of photoreceptor differentiation is simultaneously *increased* by rising *T* levels and *decreased* by falling P levels. Outside of the saturation regime— i.e. where 𝑇 < 𝑘𝑆𝑎𝑡_𝑆_, 𝑘𝑆𝑎𝑡_𝐿_— these changes precisely cancel, leaving the rate of photoreceptor differentiation unchanged. As a result, these rates do not change with *P* abundance, creating the (effectively) zero-order relationship required for noise suppression.

#### 3.5. A note on non-cone fates

An interesting feature of the signaling model is that the ‘alternative’ non-cone fate plays an integral role to the noise filtering described above. In order for the signaling domain to effectively reject noise in *P* abundance, there must be a pathway of *P* consumption with a first-order (i.e. *T*-independent) rate constant. This pathway is therefore not a ‘catch all’ for other fates accessible to *P* cells, but rather an important part of T3-regulated photoreceptor development. We also note that the identity of this ‘alternative’ fate does not matter for this role. Indeed, even cell death could serve this purpose. In this case, the noise in initial *P* numbers would be ‘absorbed’ by the dying cell population so that photoreceptor specification can be precise. In this sense, cell death could be regarded not as a wasteful side product of development, but rather as the cost of precision.

